# Improved prediction of MHC II antigen presentation through integration and motif deconvolution of mass spectrometry MHC eluted ligand data

**DOI:** 10.1101/799882

**Authors:** Birkir Reynisson, Carolina Barra, Saghar Kaabinejadian, William H Hildebrand, Bjoern Peters, Morten Nielsen

## Abstract

Major Histocompatibility Complex II (MHC II) molecules play a vital role in the onset and control of cellular immunity. In a highly selective process, MHC II presents peptides derived from exogenous antigens on the surface of antigen-presenting cells for T cell scrutiny. Understanding the rules defining this presentation holds critical insights into the regulation and potential manipulation of the cellular immune system. Here, we apply the NNAlign_MA machine learning framework to analyse and integrate large-scale eluted MHC II ligand mass spectrometry (MS) data sets to advance prediction of CD4+ epitopes. NNAlign_MA allows integration of mixed data types, handling ligands with multiple potential allele annotations, encoding of ligand context, leveraging information between data sets, and has pan-specific power allowing accurate predictions outside the set of molecules included in the training data. Applying this framework, we identified accurate binding motifs of more than 50 MHC class II molecules described by MS data, particularly expanding coverage for DP and DQ beyond that obtained using current MS motif deconvolution techniques. Further, in large-scale benchmarking, the final model termed NetMHCIIpan-4.0, demonstrated improved performance beyond current state-of-the-art predictors for ligand and CD4+ T cell epitope prediction. These results suggest NNAlign_MA and NetMHCIIpan-4.0 are powerful tools for analysis of immunopeptidome MS data, prediction of T cell epitopes and development of personalized immunotherapies.

## Introduction

Major Histocompatibility Complex class II (MHC II) molecules play a pivotal role in the adaptive immune system. Antigen-Presenting Cells (APCs) display MHC II molecules in complex with peptides^1^ on their surface. These peptides are products of extracellular proteins internalized by APCs and proteolytically digested in endocytic compartments. During protein degradation, the MHC II binding cleft is loaded with peptide fragments of the antigen and peptides that bind stably (forming a pMHCII complex) are shuttled to the cell surface for presentation to T-helper (Th) cells of the immune system^2^. Th cells scrutinize the surface of APCs and if the T cell receptor (TCR) recognizes a pMHCII complex, Th cells can become activated. Peptides that cause T cell activation are termed T cell epitopes. Regulation of Th cell activation is critical since they coordinate the activation of effector cells. Peptide-MHC presentation is a necessary and highly selective step in the process of T cell activation and characterizing the rules defining this presentation is pivotal for our understanding of cellular immunity.

Given this, characterizing the rules of MHC II peptide presentation has been the focus of considerable research efforts. MHC II is a heterodimer, the alpha and beta chains of which together form the peptide-binding cleft. In humans, the Human Leukocyte Antigen (HLA) of MHC class II is encoded by three different loci (HLA-DR, -DQ and -DP)^3^. The corresponding HLA genes have numerous allelic variants with polymorphisms that are mostly clustered around the residue locations forming the peptide binding cleft, resulting in a wide range of distinct peptide binding specificities. MHC II has an open binding cleft, allowing it to interact with peptides of a broad length range, most commonly of 13-25 amino acids^4^. The binding cleft of MHC II interacts predominantly with a 9-mer register of the interacting peptide, termed the binding core^5^. The placement of this binding core varies between peptides, resulting in peptide flanking residues (PFRs) of differing length protruding out from the binding cleft of the MHC II molecule. PFRs influence both the interaction with MHC II and the activation of T cells^6^. These facts together makes the study and prediction of MHC II peptide interaction highly challenging.

Traditionally, in vitro peptide-MHC Binding Affinity (BA) assays have been used to generate data to characterize the specificity of MHC II molecules^7^, and a range of machine-learning prediction models have been developed from this data to identify the rules of peptide-MHC binding (reviewed in^8,9^). However, evidence suggests peptide-MHCII binding affinity to be a relatively weak correlate of MHC antigen presentation^10^ and peptide immunogenicity^11^. Likewise, several studies have demonstrated that MHC-II peptide binding prediction models can benefit from being trained on so-called immunopeptidome data obtained by liquid chromatography coupled mass spectrometry(LC-MS/MS) (reviewed in^8,12^). In a typical MHC II immunopeptidome Eluted Ligand (EL) assay, MHC molecules are immunopurified from lysed antigen-presenting cells. The bound peptide ligands are next chromatographically eluted from MHC molecules and sequenced via MS/MS^13,14^. The result of such an assay is a list of peptide sequences restricted to at least one of the MHC II molecules expressed by the interrogated cell line.

The biological relevance of EL data is a major advantage over BA data. EL data implicitly contains signals from steps of MHC II antigen presentation, such as antigen digestion, MHC loading of ligands and cell surface transport. Amino acid preferences in termini of immunopeptidome ligands show clear evidence of proteolytic digestion and ligand prediction has been improved by explicitly encoding this signal in prediction models^15,16,17^. Models trained on EL data can learn the ligand length preference of MHC molecules, since this preference is inherent to EL data, but not to BA data (examples include^15,17,18^).

While most prediction methods are trained either on binding affinity (BA) or MHC MS eluted ligand (EL) data, the work by Jurtz et al.^19^ proposed an architecture where the two data types were effectively integrated into one machine learning framework. This framework has later with high success been applied to train models with state-of-the-art performance for prediction of MHC class I antigen presentation and CD8 T cell epitopes^19,20^. Recent framework refinements also cover MHC class II data, and demonstrate how integration of antigen processing signals contained within and flanking the MS EL data boosts predictive power^15^.

The immense polymorphism of MHC combined with the high experimental cost burden associated with characterizing the specificity of individual MHC molecules makes specificity characterization of all MHC molecules a prohibitively expensive undertaking. Given this, a proposed solution is pan-specific prediction models, which are trained on peptide interaction data covering large and diverse sets of MHCs with the purpose of and learning the associations between the MHC protein sequence and its peptide specificity^21,22^. The value of these models lies in their ability to predict peptide-MHC binding for all alleles of known sequence, including those characterized by limited or even no peptide-binding data^23^.

Ligands eluted from cell lines expressing only one MHC molecule can be unambiguously annotated to that allele and are termed single allele (SA) ligands. Such data is generated from genetically engineered cell lines^24,25^ or careful experimental design, i.e. matching an immunopurification antibody specific to an MHC loci with a cell line homozygous to said loci (as done most often when conducting an HLA-DR specific LC-MS/MS experiment). However, this scenario is the exception in immunopeptidomics studies. EL data from patient samples will more often be composed of peptides of mixed specificities corresponding to the different MHC molecules expressed in the given antigen-presenting cell. Such data is termed multi allele (MA) ligands and annotating MA ligands to their respective MHC in such data is not a trivial task. Ligands from MA data can be deconvoluted into separate motifs in an unsupervised manner with tools like GibbsCluster^26^ and MixMHCp^27^, but this still leaves the task of assigning motifs to their respective alleles^12^.

We recently proposed a solution to this critical challenge of MA data by an extension of the NNAlign algorithm termed NNAlign_MA^28^. NNAlign_MA handles MA data naturally by annotating MA data during training in a semi-supervised manner based on MHC co-occurrence, MHC exclusion, and pan-specific binding prediction. This algorithm is building on similar principles as the MixMHCp^27^ and MoDec^17^ methods described earlier however, with the important difference that NNAlign_MA allows to leverage information between data sets enabling accurate motif deconvolution also for data sets and alleles with limited ligand coverage (such as HLA-C for class I^28^). In the work by Alvarez et al., the ability of NNAlign_MA to automatically deconvolute and learn binding motifs from complex EL MA data was showcased in several scenarios, and the algorithm was demonstrated to have a high potential for deconvoluting MA data and to construct accurate pan-specific predictors for MHC antigen presentation. For MHC II, this early work was conducted on a small MA data set covering mainly HLA-DR alleles, and also the NNAlign_MA modelling framework used did not incorporate signals of MHC II antigen processing. Here, we extend this work adapting NNAlign_MA to allow explicit encoding of ligand context to learn signals of antigen processing. Next, we apply the method to perform motif deconvolution and train a pan-specific prediction model of MHC II antigen presentation directly from an extensive data set of MA data covering HLA-DR, DQ and DP. Benefitting from the unique abilities of NNAlign_MA to integrate mixed data types, handling ligands with multiple potential allele annotation, encode signals of antigen processing, leverage information between data sets, we investigate its power for motif deconvolution in particular for DQ and DP molecules covered with few ligands in individual MA data sets. Also, the resulting pan-specific predictor, termed NetMHCIIpan-4.0, is benchmarked against current state-of-the-art methods in terms of performance for ligand and CD4+ T cell epitope prediction.

## Materials and Methods

### Binding affinity data

Binding affinity data was gathered from the NetMHCIIpan-3.2 publication^29^. In line with the EL data set, only BA peptides of length 13-21 were included in the analysis. After filtering, the data set contained 131,077 peptides covering 59 HLA-II molecules. Peptide binding affinity data is quantitative and the measured binding affinity was transformed to fall in the range [0,1] as described in^30^.

### Eluted ligand data - Extraction

All MHC ligands in the IEDB^31^ were downloaded (January 28th 2019th) and this table served as a guide to identify publications with MHC II EL data. Publications with full HLA class II typing and more than 1000 ligands were selected for data extraction. Ligand data was subsequently extracted from the individual publications^32,33,34,35,36,37,38,39,40,41^. Supplementary materials of the publications were processed and ligands extracted along with their source proteins and associated MHC molecules. Some additional data sets were found via literature search ^25,17,42,43,44,45^ and a small in-house database of HLA-DR EL data obtained from homozygous cell lines. Only cell lines with more than 250 measured ligands were included in the final data set.

The resulting data sets consist of tables of ligands, their source proteins and a list of potential interacting MHC II molecules (the MHC molecules expressed in the given cell line). In cases of DQ or DP heterozygosity, all possible alpha and beta chain combinations were included in the list of possible MHC II molecules. All post-translationally modified peptides were excluded and only ligands of length 13-21 were included in the analysis (excluding 16.4% of the total extracted EL data), resulting in a total of 133 data sets, covering 372,639 MHC measurements, and 74 distinct MHC II alleles (3 DPA, 12 DPB, DQA 12, DQB 12, 32 DRB and 3 H-2). For a complete overview of the data, refer to Supplementary Table 1.

### Eluted ligand data - Negative peptide generation

Eluted ligand data contain only positive examples of MHC ligands. Negative data were defined as described earlier^15,20^ by randomly sampling peptides from the UniProt database^46^. Negatives were generated per data set and follow a uniform length distribution in the range 13-21. For each length, the number of negatives was equal to 5 times the number of ligands at the maximum of the ligand length distribution.

### Eluted ligand data - Context

Terminal regions inside and outside of ligands were encoded to capture signals of proteolytic digestion, as described previously^15^. A total of 12 residues were encoded for each ligand: six PFRs (three residues from the N-terminal and three from the C-terminal) and six ligand context residues from the source protein sequence (three residues upstream the ligand N-terminus and three residues downstream the C-terminus). Roughly 1% of ligands could not be mapped to a source protein, resulting in a context encoding of X’s. BA data also received a context of X’s. Negative EL data in training sets were also assigned context from their sampled source protein.

### Data set partitioning

NNAlign_MA was trained in a 5-fold Cross-Validation manner. Data was partitioned via common motif clustering^47^ to ensure that no partition shared 9-mer subsequences. BA and EL data were clustered simultaneously and next separated, resulting in a total of 10 partitions, 5 for EL data and 5 for BA data.

### Epitopes

T Cell epitopes were downloaded from the IEDB (downloaded: July 1st 2019), and filtered to include only positive MHC II peptide epitopes of length 13-21. Furthermore, epitopes measured by intracellular cytokine staining (ICS) or Tetramer/Multimer assays were included in the benchmark, resulting in a total of 1469 epitopes.

The F-rank evaluates the ability of prediction methods to prioritise epitope discovery. In short, epitopes are ranked by prediction scores amongst their length matched peptides from the antigen, and the F-rank score is the percentage of false positives, i.e. peptides with prediction scores higher than measured epitopes. An alternative F-rank scoring scheme incorporates the effects of antigen processing. Here, each antigen was *in silico* digested into k-mers (peptides of length 13-17 amino acids) and scores predicted for each. Next, epitope length matched peptides from an antigen were assigned a score by summing prediction scores of all k-mers whose predicted binding core overlap completely with the given peptide. To remove noise in the data set, and filter out potential false positive epitopes, the data set was filtered to only include epitope with an F-rank values 10.0 for at least one of the methods benchmarked.

### NNAlign_MA training

The complete model consisted of an ensemble of 150 networks with 20, 40 and 60 hidden neurons with 10 random weight initializations for each of the 5 cross-validation folds (3 architectures, 10 seeds and 5 folds). All models were trained using backpropagation with stochastic gradient descent, for 400 epochs, without early stopping, and a constant learning rate of 0.05. Only SA data was included in training for a burn-in period of 20 epochs. Subsequent training cycles included MA data.

### Sequence logos

Sequence Logos were generated with Seq2Logo^48^. The amino acid background frequency for all logos was constructed from the set of ligand source proteins. MHC II binding motifs detected in EL data sets were generated from ligand binding registers as predicted by NNAlign_MA, using the default Seq2Logo settings. For context, separate logos were made for the N- and C-terminal using weighted Kullback-Leibler, excluding sequence weighting. Logos from GibbsCluster deconvolution of context were generated with default settings of Seq2Logo. Note, that a condition for inclusion in context motif visualization was that the PFR of the ligand be 3 residues or longer, in agreement with earlier work^15^.

### Allele sequence distance measure

Sequence distance between alleles is calculated based on the alignment score of their pseudo sequences with the following relation: 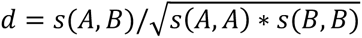, where *s*(*A*, *B*) represents the BLOSUM50 alignment score of the two pseudo sequences^49^.

### Deconvolution consistency score

As a measure of NNAlign_MA’s consistency in MHC motif deconvolution, the motifs for a given MHC as obtained from different MA data sets were compared in terms of the Pearson correlation coefficient of their PSSMs. We defined the consistency score for an allele’s deconvolution as the average Pearson correlation coefficient for non-self comparison.

## Results

The goal of this project is to build a pan-specific predictor of MHC II antigen presentation trained on EL data. Our approach is to train a model with NNAlign_MA^28^ adapted to MHC II EL data by allowing for explicit encoding of ligand context. In large-scale benchmarks, we investigate the ability of NNAlign_MA to consistently deconvolute motifs from data sets of mixed DR, DP and DQ peptide specificities. Furthermore, investigating the ligand context, we validate earlier findings demonstrating an improved predictive performance for ligand discovery empowered by the signal of antigen processing, and investigate this signal’s consistency across MHC loci. Finally, independent CD4+ epitope benchmarks compare the developed model to recent, competing models.

### NNAlign_MA algorithm

MA data are numerous and diverse, ∼73% of ligands gathered for this project stem from MA data sets (see Supplementary Table 1 for a summary of EL data sets). When dealing with EL data from MHC II heterozygous cell lines, the ligands are of mixed specificities and it is not trivial to assign ligands to their respective restricting MHC II molecule. We have earlier proposed a simple machine learning framework, NNAlign_MA, to resolve this task^28^. Using semi-supervised learning, NNAlign_MA leverages information from SA data to annotate MA data. This is achieved with a burn-in period in which only SA data is used for training, after which MA data is introduced, annotated and used for training. The annotation is achieved by predicting in every training iteration binding to all MHC molecules assigned to the given MA data set and assigning the restriction from the highest prediction value. With the allele assignment in place, the MA data becomes equivalent to SA data and can be used for training.

An important strength of NNAlign_MA is that the MA data annotation is integral to the training process, circumventing the need for offline motif deconvolution and annotation. The fully automated process implicitly leverages data set allele overlap and exclusion principles to learn motifs and annotate ligands. An example of this functionality is shown in Figure 1, displaying the motifs identified from three MA EL data sets that pairwise share one MHC allele. This figure illustrates how the model achieves allele annotation by comparing motifs and alleles shared between multiple data sets.

**Figure 1.**
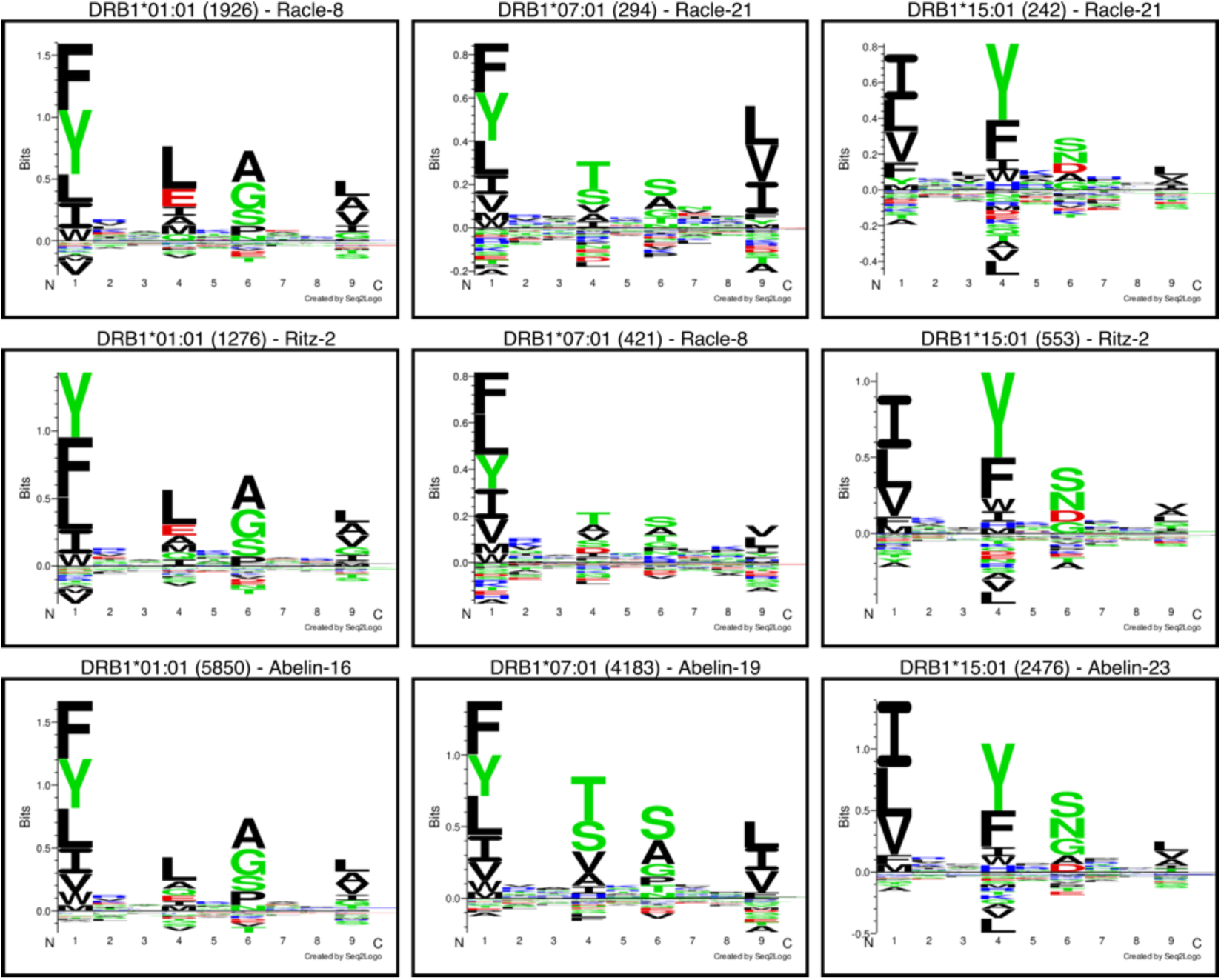
Multi-allele (MA) data motif deconvolution of 3 cell lines containing pairwise allele overlap compared to logos from 3 Single-Allele (SA) data sets. The figure shows the ability of NNAlign_MA to implicitly leverage allele overlap and the exclusion principle to annotate ligands from MA data and learn their binding motifs. The logos were generated with Seq2Logo, excluding ligands with a prediction score less than 0.01. Each logo is labelled with the allele, the number of ligands used to generate the logo and the identifier to the source data set.

### Comparison of models trained on SA and MA data

NNalign_MA extends the training data by its ability to annotate MA data automatically. To assess if NNAlign_MA can learn specificities uniquely covered by MA data without diluting learning from SA data, we compared three models in terms of cross-validation performance. One model, SA-model, is trained exclusively on SA data (SA EL data and BA data) and another, MA-model, is trained on all data (SA EL, MA EL and BA data). The third model, NetMHCIIpan-3.2, is included as a representative model trained only on BA data. Figure 2 displays the performance of the different models on the two EL SA and MA data subsets (the performance of the SA and MA models evaluated using cross-validation).

**Figure 2.**
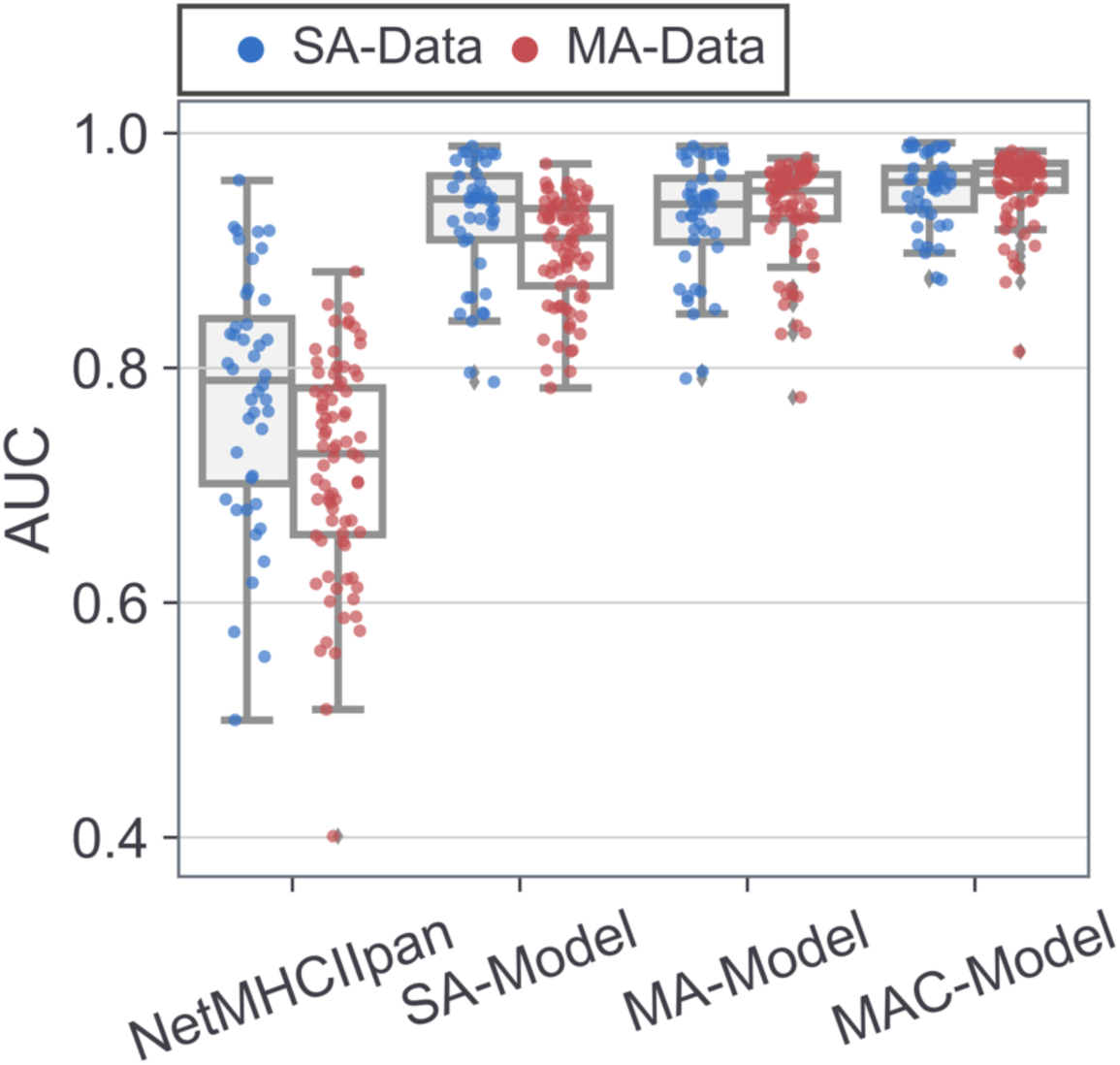
Cross-validation benchmark performance comparison of models with increasing complexity in terms of training data and encoding. SA-model and MA-model are trained on SA-data and MA data, respectively. MAC-Model is trained on MA-data, including ligand context encoding. SA-, MA- and MAC-models were evaluated on the complete data set by predicting each of the 5 test sets in a cross-validation setup, concatenating each test set result and calculating data set wise AUC scores. The four models are compared separately for SA (blue dots, n=44) and MA (red dots, n=79) EL data sets. Every point in the figure represents an AUC value for a single ligand data set. Results of binomial statistical tests, excluding ties: SA-Data comparing MA-Model and SA-Model, p-value=1.00. MA-Data comparing MA-Model and SA-Model, p-value<<0.001. All data comparing MA-Model to NetMHCIIpan-3.2, p-value<<0.001. All data comparing MA-Model and MAC-Model, p-value<<0.001.

These results demonstrate that the SA and MA models shared comparable predictive performance when evaluated on SA data (p=1.000, binomial test excluding ties), supporting that NNAlign_MA maintains performance on SA data when including MA data in training. Moreover, the MA-model significantly outperformed the SA-model when evaluated on MA-data (p-value < 10^-24^, binomial test excluding ties). This gain in performance of the MA over the SA-model correlated strongly with the average allele distance of the MA data sets to the alleles covered by SA-data (PCC: 0.732, Supplementary Figure 2). For details on this distance measure, refer to materials and methods. That is, the performance gain of the MA-model is - as expected - most pronounced for MA data sets with alleles distant to the SA data. Finally, are both the SA and the MA models demonstrated to significantly outperform NetMNHCpan-3.2 on both EL data sets (p<0.001 in all cases). Taken together, these results demonstrate that NNAlign_MA can successfully deconvolute motifs from MA data and use this to boost the predictive power beyond that of methods trained on SA data. Likewise, these results confirm the earlier finding that methods trained on EL data outperform methods trained on BA data for the task of predicting MHC eluted ligands^15^.

### Source protein context boosts ligand prediction

Immunopeptidome data inherently contains signals from steps leading to antigen presentation and patterns of proteolytic digestion have been described in the terminal and context regions of ligands^15^. Earlier work has suggested that models encoding this context information have superior performance when predicting ligand data^15,16,17^. Here, we set out to validate this observation on an extensive data set.

The encoding of context data was performed as previously described^15^ (for details refer to Materials and Methods), and models with and without encoding of ligand context were trained and evaluated using cross-validation. The results of the benchmark are shown in Figure 2 and demonstrate a significantly improved performance of the model, MAC-Model, trained including the ligand context (p<0.001 for both SA- and MA-data, binomial test excluding ties). This result thus confirms the earlier finding conducted on a data set covering only a limited set of HLA-DR alleles^15^.

### Consistent motif deconvolution from MA data

The posterior analyses, demonstrated the high predictive power of NNAlign_MA and suggested that accurate deconvolution of binding motifs in MA data is driving this performance. To further support this claim, we show in Supplementary Figure 1 binding motifs generated from the predicted binding cores of ligands in each MA data set when deconvoluted by NNAlign_MA. By visual inspection, this figure confirmed that for a vast majority of cases, NNAlign_MA was able to successfully and consistently deconvolute motifs from MA data. For most data sets, motifs captured sharp information enrichment at well-defined anchor positions (P1, P4, P6 and P9 most molecules).

A more quantitative analysis of deconvolution consistency was achieved by calculating pairwise correlations between PSSMs representing the motifs for MHC molecules shared between multiple data sets. That is, for every MHC molecule shared between 2 or more data sets, PSSMs were generated from the binding cores of ligands assigned to said allele for each data set, and the PSSMs pairwise compared by Pearson correlation. Supplementary Figure 3. visualizes this analysis in the form of heatmaps. From these heatmaps, a consistency score was calculated for a given allele from the average of these inter data set correlation values. The result of this (Supplementary Figure 4) demonstrated very high consistency values for most DR, DP and H-2 alleles (median for each locus over 0.887), and overall lower values for DQ (median 0.800).

To further quantify the accuracy of individual motif deconvolutions obtained by NNAlign_MA, the positive predictive value (PPV0.9) was calculated for each set of positive peptides assigned to the given MHC molecule within each data set in the context of a set of random natural negatives. In short, the PPV0.9 is calculated from the proportion of true positives within the top N predictions in the context of negative decoy peptides. Here N was defined as 90% (to account for noise in the EL data) of the number of ligands assigned to the given allele in a given data set, and the negative decoy peptides outnumbered positives 99 to 1. The result of this analysis is presented in Supplementary Table 2 and Supplementary Figure 5, and overall confirmed a high performance with median PPV0.9 values for DR, DP and H-2 alleles over 0.604. However, also here the performance for DQ was found to be reduced (median PPV0.9=0.372).

Combined, these results support the claim that NNAlign_MA can accurately and consistently deconvolute binding motifs in EL MA data. We further in Figure 3. quantify the number of MHC-II molecules identified with a PPV0.9 above the conservative value of 0.3 (30% of the predicted positives are ligands, compared to the performance of 1% of a random predictor). Here NNAlign_MA could identify accurate motifs for 31 HLA-DR (one molecule, DRB1*03:05, was not captured due to only being characterized with 90 ligands falling below the threshold of 100 imposed for accurate motif characterization (see supplementary figure 1), 7 HLA-DP, 12 HLA-DQ, and 2 H-2 molecules (see Supplementary Figure 6 for an overview of the motifs for these molecules identified between different data sets). In particular for DR these numbers are impressive, with a coverage of 97% of the alleles in the training data, and these numbers largely surpass the set of molecules described in the recent Racle^17^ and Abelin^25^ publications suggesting a superior capacity for motif deconvolution of NNAlign_MA compared to the MoDec method proposed by Racle et al. in particular for molecules characterized by limited ligand annotations in individual MA data sets. The number of MHC molecules covered by accurate motif characterization however is in general lower for DQ (and DP) compared to DR reflecting the overall low number of ligands annotated to MHCs from these loci (on average less than 23% of ligands from pan-class II MA data sets are annotated to DP and DQ molecules) and the associated challenge this imposes when identifying accurate binding motifs.

**Figure 3.**
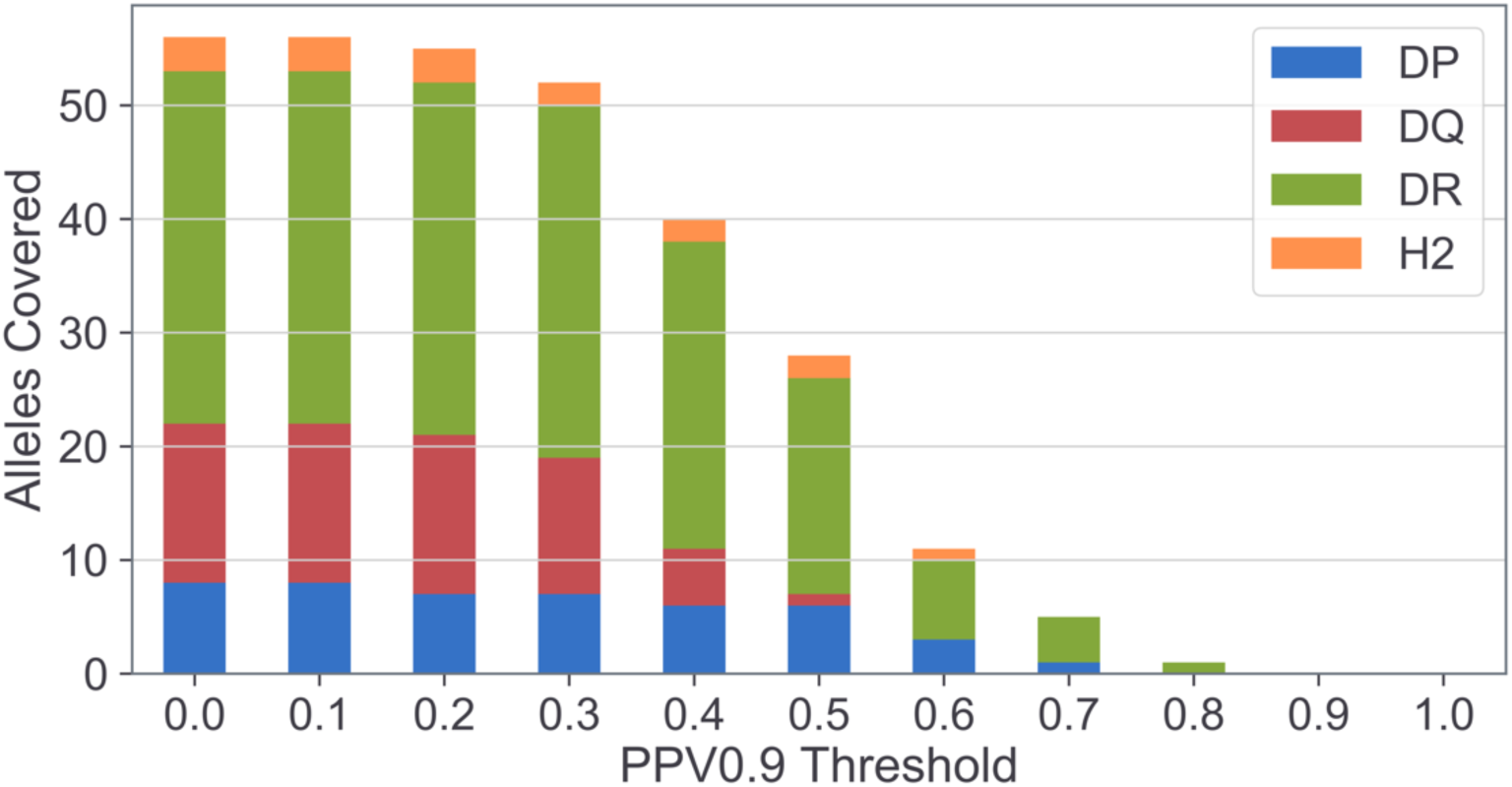
Number of MHC II molecules covered at different thresholds of PPV0.9. Each molecule is represented by the average PPV0.9, across data sets. For PPV calculations, the ligand/decoy ratio is 1/99 for a set of decoy peptides independent of training data.

### Pan-specific prediction power

A central power of the NNAlign_MA modelling framework is its ability to leverage information between data sets, boosting performance for molecules with limited ligand coverage combined with its pan-specificity allowing to make accurate predictions outside the set of molecules included in the training data. The first part was illustrated above with the expanded number of molecules characterized by accurate motifs in comparison to the method by Racle et al. While the pan-specific prediction power of the original NNAlign model has been documented previously using different kinds of leave-one-allele-out experiments^18,21^, this is only the case to a minimal extent for NNAlign_MA^28^. The reason for this is that leave-one-out experiments cannot readily be conducted on MA data sets since molecules in this situation in general are shared between multiple data sets making it non-trivial to define what to leave-out. As an alternative, we identified 3 molecules characterized only with SA-EL data in our training data set (DRB1*04:02, H2-IAb and DQA1*01:02-DQB1*06:04). Leaving out all (both BA and SA) data for these molecules from the training data, the predictive model was retrained and the predictive performance for each molecule evaluated, resulting in an average AUC performance of 0.896 (compared with the cross-validation performance values of these data sets of 0.950). To further assess the quality of the pan-specific predictions, we in Figure 4 show the binding motifs for the three molecules as obtained from the SA data themselves, the motifs predicted by this leave-molecules-out (LMO) model, and the motifs for the corresponding nearest neighbor alleles contained within the training data set (identified using the distance measure defined in material and methods). This latter motif would mimic the performance of a the strategy of training allele specific models and using nearest neighbor inference to make predictions for alleles outside the training set. Overall, these results show a high agreement with only minor differences (including the “missed” P3 anchor for DQA1*01:02-DQB1*06:04) between the “true” motifs of the three molecules and the motifs predicted by the leave-out model, and a likewise poorer predictive power of the nearest neighbor approach in particular for the molecules with more distant nearest neighbors (nearest neighbor distances to training set: DRB1*04:02, 0.08; DQA1*01:02-DQB1*06:04, 0.11; H-2-IAb, 0.33).

**Figure 4.**
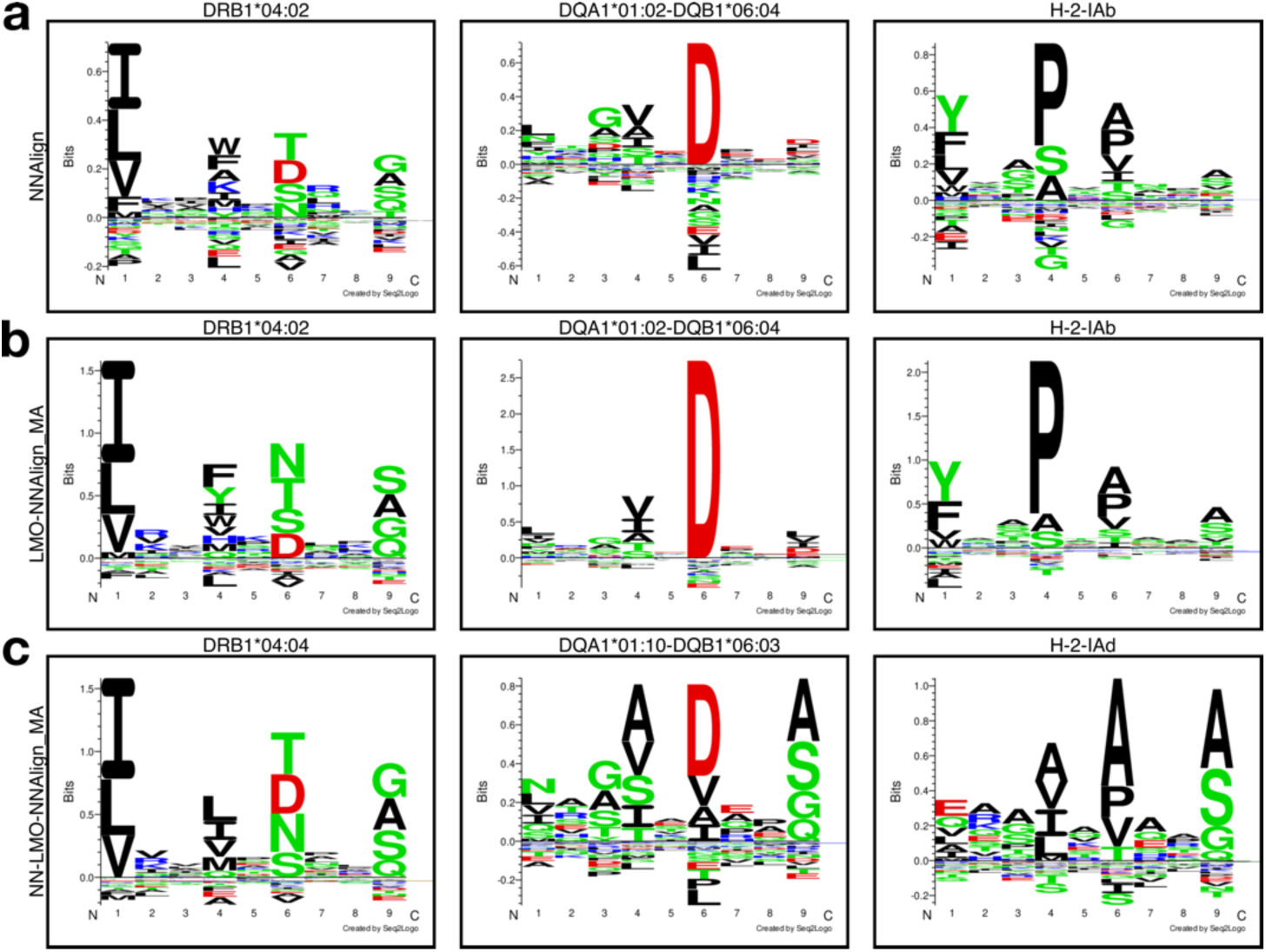
Motifs for three alleles only covered by EL SA data. **a.** Logos generated from binding cores of a model trained only on the available EL data using NNAlign^50^. **b.** The learned motifs of a model trained leaving out the three alleles in question, generated from the predicted binding cores of the top 0.1% of a set of 1,000,000 random natural 15-mer peptides. **c.** The learned motifs of the same model as in **b.** but for nearest neighbor alleles in the training set for the left out alleles, generated from the predicted binding cores of the top 0.1 % of a set 1,000,000 random natural 15-mer peptides.

### Signal of proteolytic antigen processing

We have previously demonstrated how incorporation of ligand source protein context boosts the predictive power of NNAlign_MA (Figure 2). To further investigate the source of this, and seek to relate the findings to specific proteolytic signals of antigen processing, we in Figure 5, show sequence logos representing the N and C terminal context signal contained within the EL data set. Here, all ligands were mapped to their antigen source protein to extract the ligand context of 6 residues (3 upstream and 3 downstream of the ligand). Along with the context, 3 residues at each terminus were extracted. A few observations can be made from this figure: firstly, the ligand contains more signal than the context (Figure 5a). Secondly, the data shows a pronounced enrichment of Proline in positions N+2 and C-2 in agreement with earlier findings^15^. Finally, both termini are enriched for charged amino acids. An unsupervised deconvolution of the context motif (using GibbsCluster^26^) supports the two latter observations (Figure 5b). The enrichment for P at positions N+2 and C-2 is accentuated in the deconvolution, along with other well-defined motifs. This supports the notion that proteolytic enzymes of more than one specificity are at play in the degradation of MHC II antigens, in agreement with earlier findings^16,17^. Further, evaluating the predictive power for ligands assigned to each of the different clusters revealed a consistent high predictive performance across all clusters (median values above 0.975 in all cases) and a slightly improved performance for the clusters with P at position N+2 and C-2 both for N- and C-termini (AUCs computed data set wise for each cluster, with negatives assigned to clusters proportionally to ligand cluster assignment). Analyzing the data separately for DP, DQ, DR and H2 molecules revealed very similar results (see Supplementary Figures 7 and 8), again suggesting that the observed signal and identified motifs are related to antigen processing and not MHC binding.

**Figure 5.**
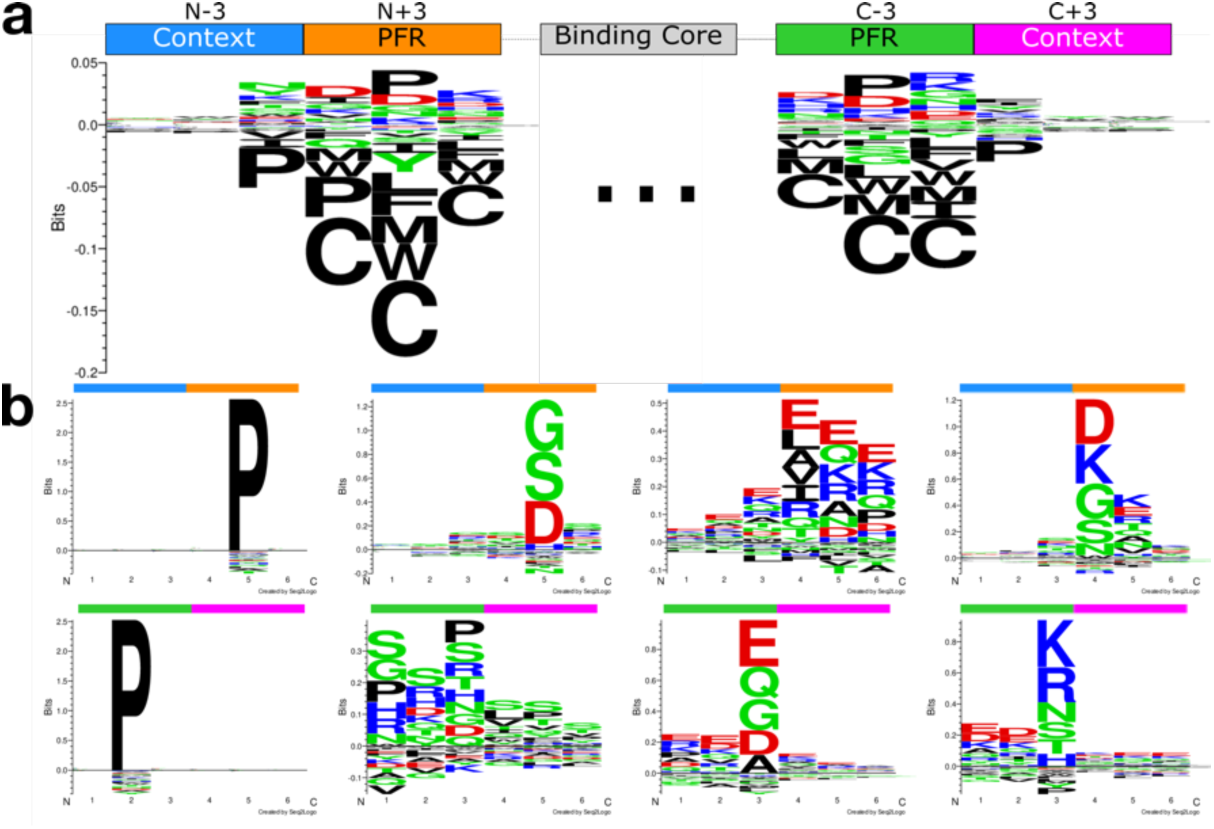
Sequence logos for N-(top left) and C-terminal (top right) and context sequences. **a.** Logos for the N- and C-terminus of all ligands (with PFRs of length >2). The diagram indicates the placement of residues relative to the N- and C-terminus of ligands. **b.** Upper and lower panel show, unsupervised GibbsCluster deconvolution of N- and C-terminal contexts, respectively. Cluster motifs have been ordered manually to align similar motifs for the N- and C-terminus, with the Proline motif having highest information content in both cases.

### CD4+ epitope evaluation

Prediction of CD4+ epitopes is the ultimate benchmark of peptide-MHC II binding predictors. Earlier work has suggested that prediction methods trained including EL data share improved predictive performance for this task compared to method trained on BA data, but also that the gained performance for prediction of MS ligands observed when including context information only to a minor degree - if at all - is transferred to epitope identification^15,16^. Here, we set out to test if the models developed here align with these findings. That is, we benchmarked the predictive performance of models trained with (MAC-model) and without (MA-Model) context encoding on a large set epitopes from the IEDB and compared performance to the state of the art methods NetMHCIIpan-3.2^29^ (trained on BA data only), MHCnuggets^51^ (trained on BA and EL data), and MixMHCIIpred^17^ (trained on EL data). Note that other methods for predicting MHC II binding trained on EL data have been proposed, including NeonMHC2^25^ and MARIA^52^. However, the NeonMHC2 only allows one to run max 20 predictions per day, and MARIA only 5000 predictions per submission making it impractical to include in a benchmark covering more than 700,000 peptide-MHC combinations. Benchmark performance was evaluated using the F-rank score. In short, F-rank is the proportion of false-positive predictions within a given epitope source protein, i.e. percentage of peptides with a prediction score higher than that of the epitope. An F-rank value of 0 corresponds to a perfect prediction (the known epitope is identified with the highest predicted binding value among all peptides found within the source protein) and a value of 50.0 to random prediction. Figure 6 gives the results of this experiment showing F-rank values obtained by *in silico* digesting the epitope source protein into overlapping peptides of the length of the epitopes. The benchmark is split into two subsets, one with (843 epitopes covering 20 MHC II molecules) and one without the HLA-II molecules covered by MixMHC2pred (149 epitopes covering 16 MHC II molecules). This benchmark demonstrated a significantly improved performance of the MA-model compared to NetMHCIIpan-3.2, MixMHC2pred, MHCnuggets and the MAC-Model (p<0.028, p<<0.0001, p<<0.0001 and p<<0.0001, respectively for a binomial test excluding ties). Inspecting further epitopes of alleles covered by MixMHC2pred showed that NNAlign_MA’s gain in performance was not only driven by epitopes of a limited set of restrictions, but hold generally across the vast majority of HLA molecule included in the benchmark (improved median F-rank for 16 out of 20 restrictions, 1 tie, p-value<0.005 in binomial test, Supplementary Figure 9). Likewise, the MA-model showed significant improvement compared to NetMHCIIpan-3.2 and MHCnuggets for the subset of epitopes with MHC restriction covered by the training data (p<0.012 and p<<0.0001, respectively, binomial test excluding ties). This gain in performance is further maintained when analyzing the subset of epitope restricted to MHC molecules not included in the training data, again highlighting the pan-specific prediction power of the NNAlign_MA modelling framework.

**Figure 6.**
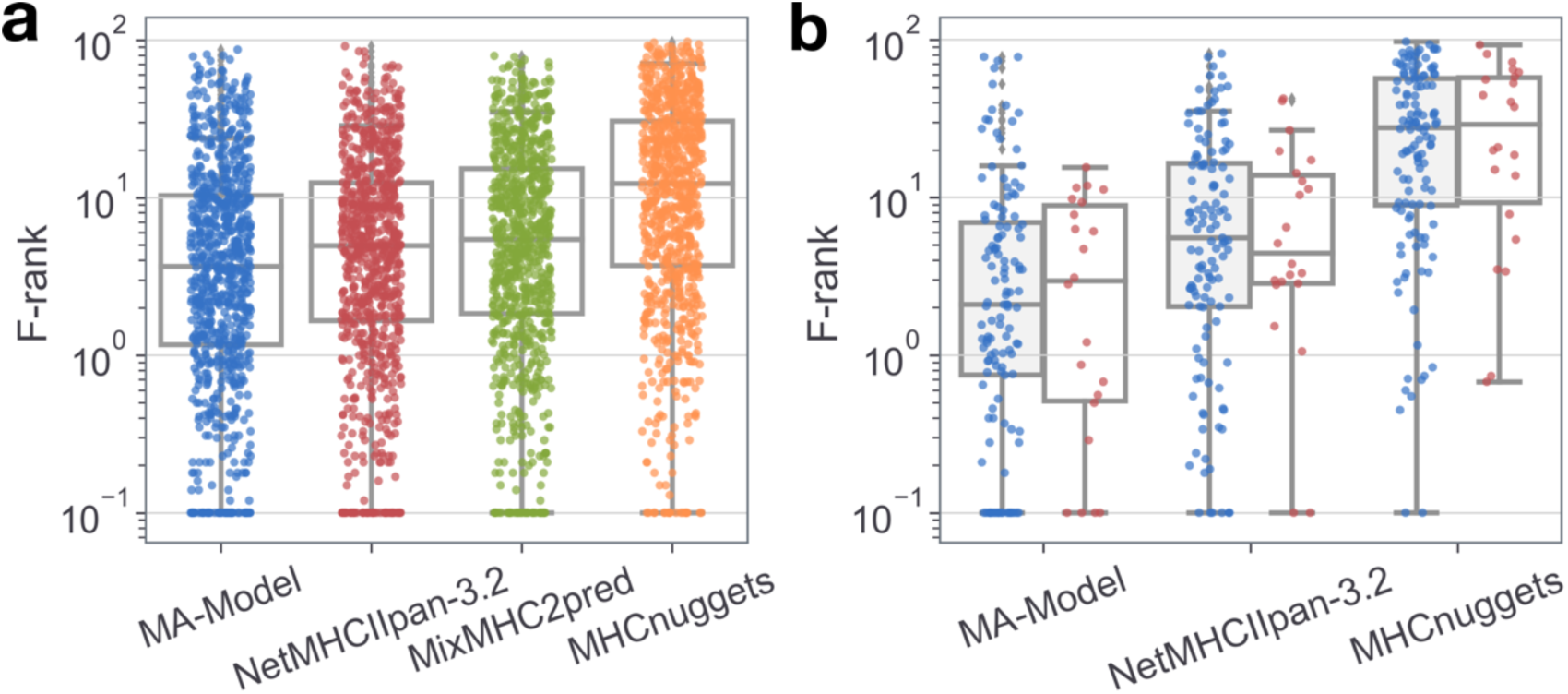
F-rank evaluation of NNAlign_MA, NetMHCIIpan-3.2 and MixMHC2pred models on ICS+Tetramer epitopes from the IEDB. **a.** MA-Model (blue) F-rank is compared to NetMHCIIpan-3.2 (red), MixMHC2pred (green) and MHCnuggets (orange) for epitopes of alleles covered by the MixMHC2pred method (843 epitopes). **b.** Epitope F-rank of MA-Model, NetMHCIIpan-3.2 and MHCnuggets are compared for epitopes of alleles covered by the MA EL training data (blue) (127 epitopes) and epitopes of alleles outside the training data (red) (22 epitopes). **a and b**: Each point in the plot represents the F-rank of one epitope, the F-rank being the percentage of false-positive length matched peptides from the epitope source protein, as described in methods. Only results for epitopes for which at least one prediction method had F-rank below 10.0 are presented. For visualization, F-rank values of 0 are presented as 0.1005.

The results however also confirm the earlier finding that signals of antigen processing contained within peptide context did not benefit the predictive power for epitope discovery. As discussed earlier^15,16^, this observation might not come as a surprise, since peptides tested for T cell immunogenicity most commonly are generated as overlapping peptides spanning a source protein, and hence are not expected to follow any rules of antigen processing. However, even attempting to account for this bias applying a scoring scheme where the score of a given peptide was assigned from the sum of the individual prediction values from all 13-17-mer peptides with predicted binding core overlapping the given peptide (for details see Materials and Methods) maintained the improved predictive performance of the MA-model and hence did not alter this conclusion.

These findings thus align with earlier results^15,16^ demonstrating an improved performance for CD4+ epitope identification of models trained including EL data compared to methods trained on binding affinity data only (NetMHCIIpan-3.2), and that adding context information to such predictors does not boost performance for prediction of epitopes despite this being observed for prediction of MHC ligands.

### Neoepitope evaluation

A final benchmark compared the performance of NNAlign_MA and MixMHC2pred for prediction of CD4+ neoepitopes. A recent study^17^ compiled CD4+ neoepitope immunogenicity measures from patients with full DP, DQ and DR typing, including both positive and negative results. Here, we present a comparison of prediction performance on this epitope set in terms of AUC and AUC0.1 (Figure 7). Excluding peptides with a length less that 13 amino acids resulted in a benchmark of 928 peptides. For predictions, to make a fair comparison between the two methods, only HLA molecule covered by MixMHC2pred were included. MixMHC2pred prediction were performed as described in^17^ by assigning a prediction value to each peptides as the high prediction score of all overlapping 15-mers to all HLA molecules of the patient covered by MixMHC2pred. For NNAlign_MA, the sub-kmer approach described earlier^16^ was applied. Here predictions were made for all 13-21-mers subsequences contained within a given peptide to all HLA molecules of the patient covered by MixMHC2pred. Next, prediction scores were transformed into percentile ranks by reference to a prediction score distribution generated from 100,000 random natural peptides and a score was assigned to each kmer as the lowest percentile rank across all HLAs. Finally, a score to each peptides was assigned as the average over all k-mers. The result of this analysis demonstrated a slightly higher AUC of NNAlign_MA compared to MixMHC2pred. However, focusing on the earlier part of the ROC curve, the difference becomes more substantial with AUC0.1 values of 0.273 (NNAlign_MA) and 0.182 (MixMHC2pred), suggesting that NNAlign_MA reliably can identify a larger proportion the of the neoepitopes. By way of example, the two methods identify 44% (NNAlign_MA) and 37% (MixMHC2pred) of the neoepitopes at a false positive rate of 10%.

**Figure 7.**
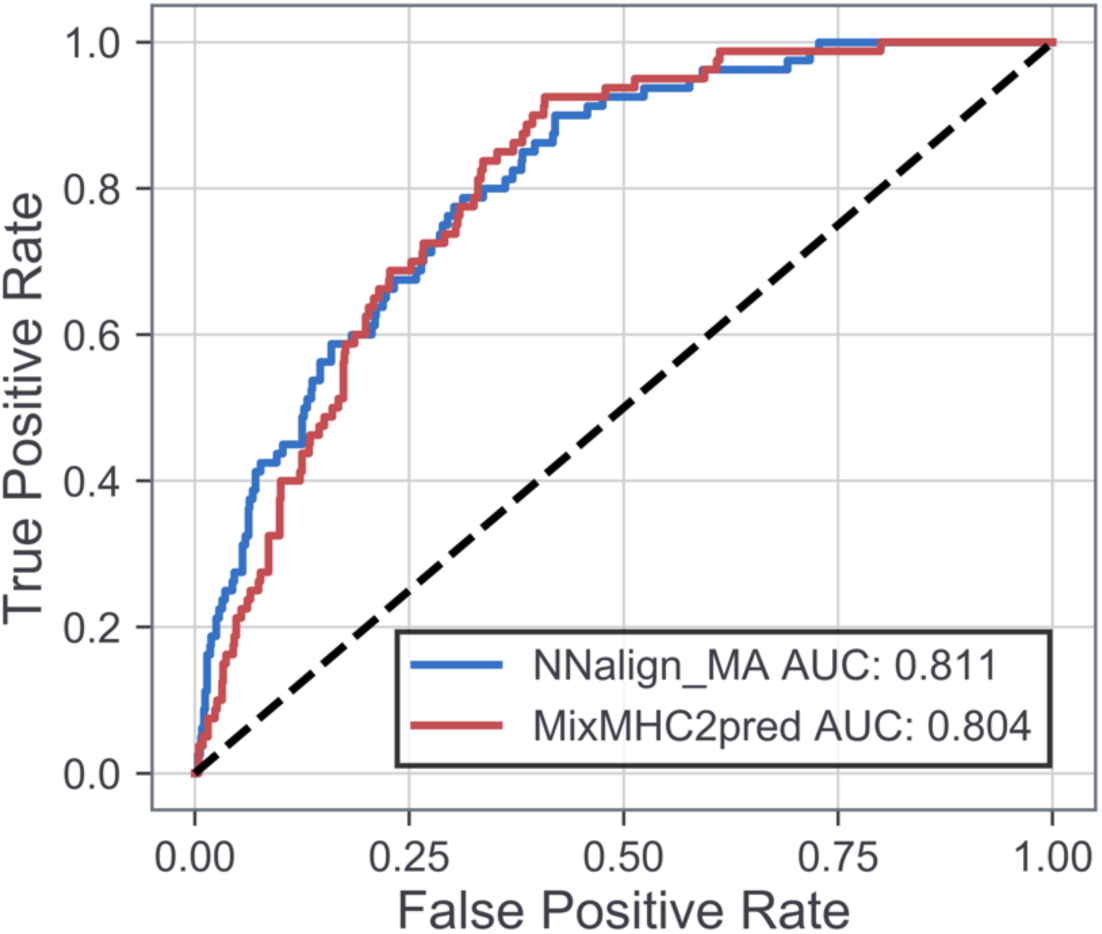
Neoepitope benchmark comparing NNAlign_MA and MixMHC2pred. The receiver operator curve for the prediction of a CD4+ neoepitope benchmark data set (n=928).

### NetMHCIIpan-4.0

A web server implementation of the NNAlign_MA and NNAlign_MAC models is available at http://www.cbs.dtu.dk/services/NetMHCIIpan-4.0/index_v0.php

## Discussion

In this work, we have trained and evaluated a pan-specific MHC II binding predictor. This predictor integrates recent advances towards full utilization of EL data for prediction of MHC antigen presentation: two output neuron architecture allowing integration of multiple data types, the encoding of proteolytic ligand context, and use of the NNAlign_MA machine learning framework allowing training on multi-allele (MA) EL data. Training on an extensive EL data set, we have in a stepwise manner shown how these features come together to construct a prediction model that significantly improves upon current state-of-the-art methods, including its predecessor NetMHCIIpan-3.2 when it comes to predicting both ligands and CD4+ epitopes.

Firstly, we provided examples of how NNAlign_MA consistently deconvolutes MHC binding motifs from separate MA data sets confirming that NNAlign_MA provides accurate solutions to the challenge of annotating binding motifs and learning from MA data.

Secondly, MA-models outperformed both SA-models and NetMHCIIpan in cross-validation evaluation of MA data, and MA models and SA models had comparable performance on SA data supporting the claim that MA data did not dilute the learning and performance of NNAlign_MA. Relatedly, the largest gain of MA over SA models was for data sets that contained alleles distant to alleles covered by SA data. This result underlines how NNAlign_MA successfully increases the coverage of predictable alleles by deconvoluting binding motifs in and training on MA data. This observation is critical for our goal of training a pan-specific MHC II predictor on EL data, as the success of pan-specific predictors is contingent on training data with broad allelic coverage^21,53^.

Thirdly and in line with earlier work^15,17^, encoding proteolytic context in models improved performance for prediction of ligands significantly. We found that proteolytic context of ligands is a general feature that holds across MHC II loci and species. Clustering context information resulted in motifs with a strong signal of amino acid enrichment/depletion in the peptide flanking regions. These motifs are likely a signature of proteolytic cleavage processes taking place in the MHC II antigen presentation pathway^16^. Across loci and species, we observed a motif of Proline enrichment in the N+2 and C-2 positions of ligands. This is consistent with the hypothesis of blocked endopeptidase digestion by the secondary α-amine of Proline. This Proline enrichment has been associated with peptides processed in the endo-lysosomal pathway of MHC II antigen presentation^54^. Likewise, we consistently observed motifs with enrichment for aspartic and glutamic acid. We suggest this enrichment reflects Cathepsin disfavoring negatively charged amino acids near cleavage sites^55^, leading to decreased digestion of ligands with said signature. With lesser information content, a motif of the N-terminal context deconvolution displayed enrichment of glycine, serine and threonine. A similar enrichment has been observed downstream of the cleavage site for Cathepsin S, a protease known to participate in the MHC II antigen presentation pathway^56^. The identified processing motifs are thus to a high degree associated with decreased potential for antigen processing. Given this, we speculate that they could serve as novel MHC II agnostic ways for engineering enhanced/reduced peptide and protein immunogenicity; that is amino acid mutations away from these motifs would lead to enhanced potential for cleavage, leading to a lower level of antigen presentation, and hence a reduction in potential immunogenicity. Likewise, could mutations into these motifs, lower the potential for antigen processing, inducing increased immunogenicity.

The proposed model improved CD4+ epitope prediction when compared to the state-of-the-art predictors NetMHCIIpan-3.2, MHCnuggets and MixMHC2pred. In contrast to the findings for MS ligands, in this benchmark, the predictive power of the models did not benefit by integration of context encoding. This was observed even when accounting for bias in the data imposed by T cell epitope peptides most often being generated synthetically and thus not reflecting rules of natural antigen processing. This result is in line with earlier observations^15,16^ and suggests that the cleavage signal, even though present in individual ligands are diluted when analyzing T cell epitopes due to the vast volume of possible ligands with binding cores overlapping the epitope.

Finally, the performance of a model for identification of CD4+ cancer neoepitopes was benchmarked against MixMHC2pred. Here, the two methods were found to have overall comparable predictive performance, with NNAlign_MA demonstrating the highest specificity and MixMHC2pred the highest sensitivity. From a clinical and applied perspective, this result suggests NNAlign_MA (and the resulting predictor NetMHCIIpan-4.0) to be a useful to tool as a guide also for identification of neoepitopes.

A key factor determining the power of a predictive model for MHC antigen presentation is the volume and MHC diversity of the training data. Here, we have demonstrated how the ability of NNAlign_MA to deconvolute motifs and integrate MA EL data into the training framework allows for an effective increase of the MHC diversity of the training data. By way of example, the training data used in the current study is limited to 22 (14 DR, 2 DP, 5 DQ, 1 H2) molecules if focusing only on SA EL data. By integrating MA data, this number is expanded to 102 (32 DR, 18 DP, 49 DQ, 3 H2) (accounting for all possible α and β chain combinations in heterozygous DQ and DP data sets). However, this HLA diversity is only relevant if associated with large volumes of peptide data. Analyzing the results of the motif deconvolution of NNAlign_MA on the training data for data sets revealed a high bias in the EL data with very limited ligands annotated to DQ (and to a lesser degree DP) molecules. This bias is more substantial than what was expected from the difference in relative expression of MHC II between the different loci and is more likely caused by differences in the specificities of the antibodies used for immunoprecipitation (IP) when purifying DR versus DQ/DP MHC complexes before running the MS. Recently, Abelin et al.^25^ have proposed a framework to experimentally generate single-allele MHC MS eluted ligand data sets to resolve this issue (and the general issues of handling MA data sets) and demonstrated how this approach could be used to generate high volumes of SA EL data also for DQ and DP molecules. Alternative approaches would include working with antibodies with improved DQ (and DP) specificities and with these conduct sequential IPs allowing for identification of larger volumes of ligands for DQ and DP.

Another complicating factor for analyzing and interpreting MA EL data for DQ and DP stems from the fact that both the alpha and beta chains are polymorphic, making it non-trivial to assess which of the four possible combinations of alpha and two beta chains are presented as HLA molecules in a given cell. However, carefully selecting homozygous cell lines or cell lines with only one expressed alpha chain could help resolve this, and in this context, EL data covering DQ and DP can without any further complication be integrated into the NNAlign_MA modelling framework.

A fundamental property of NNAlign_MA setting it apart from most other methods for motif deconvolution methods and predictors for MHC antigen presentation, is its pan-specific prediction power. As illustrated here, the power allows the method to leverage information between data sets enabling the method to identify accurate binding motifs also for molecules characterized by limited ligand counts in individual data sets. This ability allowed us to expand the coverage of MHC molecules in the training data with accurate binding motifs compared to that obtained with data set-specific motif deconvolution methods such as MoDec. Beyond identifying motifs for a higher number of MHC molecules, this pan-specific power has large impacts on the application of the prediction models developed. Here, the allele-specific models, such as MixMHC2pred developed using the MoDec deconvolution framework, has predictive power limited to the few HLA molecules with accurate motif deconvolution covered in the training data.

In contrast, the pan-specific method proposed here allows accurate predictions also outside this limited space of HLA molecules. By way of example, MixMHC2pred covers 33 HLA II molecules corresponding to cases where accurate motifs could be identified from the training data. By use of NNAlign_MA, the number of molecules with characterized motifs is increased to 52, and by expanding using the pan-specific power of the trained prediction model, the coverage is extended to 1913 (including HLA-II molecules with a pseudo-sequence distance <0.05 to any of the molecules with accurate binding motifs).

Properties other than antigen processing and HLA binding, such as protein expression, and differential access to the MHC-II presentation pathway, contribute to the likelihood of antigen presentation. In recent papers (including Abelin et al.^25^ and Chen et al.^52^) have suggested modelling frameworks integrating a panel of such properties, suggesting large improvement for prediction of HLA ligandomes. Further work remains to be done to validate the generality of these findings and their potential impact on general rational epitope discovery.

In conclusion, we have shown how the relatively simple NNAlign_MA machine learning framework can be applied to deconvolute binding motifs in MHC class II MA EL data sets, and how this deconvolution allows for identification of signals associated with antigen processing, and construction of a pan-specific prediction model with significantly improved performance compared to state of the art for prediction of both eluted ligands and CD4+ T cell epitopes.

## Supporting Information

The following supporting information is available free of charge at ACS website http://pubs.acs.org

Supplementary Figure 1) Predicted binding core motif logos deconvoluted from all MA data sets.

Supplementary Figure 2) Evidence of the benefit of extending training data scope of MA data.

Supplementary Figure 3) Quantitative analysis of MA data deconvolution consistency.

Supplementary Figure 4) Kernel density estimate plot of consistency scores, colored by loci.

Supplementary Figure 5) PPV0.9 distributions of MHC deconvolutions.

Supplementary Figure 6) MHC II logos of alleles confidently learned by the trained model.

Supplementary Figure 7) Context logos from different MHC loci: DP, DQ, DR and H2.

Supplementary Figure 8) Unsupervised deconvolution of context motifs via Gibbs clustering.

Supplementary Figure 9) F-rank results of ICS+Tetramer benchmark split up by MHC restriction.

Supplementary Table 1) Metadata for EL data sets.

Supplementary Table 2) PPV0.9 of deconvolutions.

## Notes

#### Summary of Updates

Training data extended to include data from recently published studies. More extensive analysis of MHC II binding motifs learned by the network. Validation of the pan-specific power of the method. Neoepitope benchmark to further support the broad range of applicability of the method. The text was revised extensively.

## References

1. Unanue, E. R.; Turk, V.; Neefjes, J. Variations in MHC Class II Antigen Processing and Presentation in Health and Disease. Annu. Rev. Immunol. 2016, 34, 265–297. https://doi.org/10.1146/annurev-immunol-041015-055420.

2. Wieczorek, M.; Abualrous, E. T.; Sticht, J.; Álvaro-Benito, M.; Stolzenberg, S.; Noé, F.; Freund, C. Major Histocompatibility Complex (MHC) Class I and MHC Class II Proteins: Conformational Plasticity in Antigen Presentation. Front. Immunol. 2017, 8, 292. https://doi.org/10.3389/fimmu.2017.00292.

3. Traherne, J. A. Human MHC Architecture and Evolution: Implications for Disease Association Studies. Int. J. Immunogenet. 2008, 35 (3), 179–192. https://doi.org/10.1111/j.1744-313X.2008.00765.x.

4. Chicz, R. M.; Urban, R. G.; Lane, W. S.; Gorga, J. C.; Stern, L. J.; Vignali, D. A.; Strominger, J. L. Predominant Naturally Processed Peptides Bound to HLA-DR1 Are Derived from MHC-Related Molecules and Are Heterogeneous in Size. Nature 1992, 358 (6389), 764–768. https://doi.org/10.1038/358764a0.

5. Andreatta, M.; Jurtz, V. I.; Kaever, T.; Sette, A.; Peters, B.; Nielsen, M. Machine Learning Reveals a Non-Canonical Mode of Peptide Binding to MHC Class II Molecules. Immunology 2017, 152 (2), 255–264. https://doi.org/10.1111/imm.12763.

6. Godkin, A. J.; Smith, K. J.; Willis, A.; Tejada-Simon, M. V.; Zhang, J.; Elliott, T.; Hill, A. V. Naturally Processed HLA Class II Peptides Reveal Highly Conserved Immunogenic Flanking Region Sequence Preferences That Reflect Antigen Processing Rather than Peptide-MHC Interactions. J. Immunol. Baltim. Md 1950 2001, 166 (11), 6720–6727. https://doi.org/10.4049/jimmunol.166.11.6720.

7. Buus, S.; Sette, A.; Colon, S. M.; Miles, C.; Grey, H. M. The Relation between Major Histocompatibility Complex (MHC) Restriction and the Capacity of Ia to Bind Immunogenic Peptides. Science 1987, 235 (4794), 1353–1358. https://doi.org/10.1126/science.2435001.

8. Gfeller, D.; Bassani-Sternberg, M. Predicting Antigen Presentation-What Could We Learn From a Million Peptides? Front. Immunol. 2018, 9, 1716. https://doi.org/10.3389/fimmu.2018.01716.

9. Nielsen, M.; Lund, O.; Buus, S.; Lundegaard, C. MHC Class II Epitope Predictive Algorithms. Immunology 2010, 130 (3), 319–328. https://doi.org/10.1111/j.1365-2567.2010.03268.x.

10. Bassani-Sternberg, M.; Bräunlein, E.; Klar, R.; Engleitner, T.; Sinitcyn, P.; Audehm, S.; Straub, M.; Weber, J.; Slotta-Huspenina, J.; Specht, K.;, et al. Direct Identification of Clinically Relevant Neoepitopes Presented on Native Human Melanoma Tissue by Mass Spectrometry. Nat. Commun. 2016, 7, 13404. https://doi.org/10.1038/ncomms13404.

11. Backert, L.; Kohlbacher, O. Immunoinformatics and Epitope Prediction in the Age of Genomic Medicine. Genome Med. 2015, 7, 119. https://doi.org/10.1186/s13073-015-0245-0.

12. Alvarez, B.; Barra, C.; Nielsen, M.; Andreatta, M. Computational Tools for the Identification and Interpretation of Sequence Motifs in Immunopeptidomes. Proteomics 2018, 18 (12), e1700252. https://doi.org/10.1002/pmic.201700252.

13. Dudek, N. L.; Croft, N. P.; Schittenhelm, R. B.; Ramarathinam, S. H.; Purcell, A. W. A Systems Approach to Understand Antigen Presentation and the Immune Response. Methods Mol. Biol. Clifton NJ 2016, 1394, 189–209. https://doi.org/10.1007/978-1-4939-3341-9_14.

14. Caron, E.; Kowalewski, D. J.; Chiek Koh, C.; Sturm, T.; Schuster, H.; Aebersold, R. Analysis of Major Histocompatibility Complex (MHC) Immunopeptidomes Using Mass Spectrometry. Mol. Cell. Proteomics MCP 2015, 14 (12), 3105–3117. https://doi.org/10.1074/mcp.O115.052431.

15. Barra, C.; Alvarez, B.; Paul, S.; Sette, A.; Peters, B.; Andreatta, M.; Buus, S.; Nielsen, M. Footprints of Antigen Processing Boost MHC Class II Natural Ligand Predictions. Genome Med. 2018, 10 (1), 84. https://doi.org/10.1186/s13073-018-0594-6.

16. Paul, S.; Karosiene, E.; Dhanda, S. K.; Jurtz, V.; Edwards, L.; Nielsen, M.; Sette, A.; Peters, B. Determination of a Predictive Cleavage Motif for Eluted Major Histocompatibility Complex Class II Ligands. Front. Immunol. 2018, 9, 1795. https://doi.org/10.3389/fimmu.2018.01795.

17. Racle, J.; Michaux, J.; Rockinger, G. A.; Arnaud, M.; Bobisse, S.; Chong, C.; Guillaume, P.; Coukos, G.; Harari, A.; Jandus, C.;, et al. Robust Prediction of HLA Class II Epitopes by Deep Motif Deconvolution of Immunopeptidomes. Nat. Biotechnol. 2019. https://doi.org/10.1038/s41587-019-0289-6.

18. Garde, C.; Ramarathinam, S. H.; Jappe, E. C.; Nielsen, M.; Kringelum, J. V.; Trolle, T.; Purcell, A. W. Improved Peptide-MHC Class II Interaction Prediction through Integration of Eluted Ligand and Peptide Affinity Data. Immunogenetics 2019, 71 (7), 445–454. https://doi.org/10.1007/s00251-019-01122-z.

19. Jurtz, V.; Paul, S.; Andreatta, M.; Marcatili, P.; Peters, B.; Nielsen, M. NetMHCpan-4.0: Improved Peptide-MHC Class I Interaction Predictions Integrating Eluted Ligand and Peptide Binding Affinity Data. J. Immunol. Baltim. Md 1950 2017, 199 (9), 3360–3368. https://doi.org/10.4049/jimmunol.1700893.

20. Nielsen, M.; Connelley, T.; Ternette, N. Improved Prediction of Bovine Leucocyte Antigens (BoLA) Presented Ligands by Use of Mass-Spectrometry-Determined Ligand and in Vitro Binding Data. J. Proteome Res. 2018, 17 (1), 559–567. https://doi.org/10.1021/acs.jproteome.7b00675.

21. Hoof, I.; Peters, B.; Sidney, J.; Pedersen, L. E.; Sette, A.; Lund, O.; Buus, S.; Nielsen, M. NetMHCpan, a Method for MHC Class I Binding Prediction beyond Humans. Immunogenetics 2009, 61 (1), 1–13. https://doi.org/10.1007/s00251-008-0341-z.

22. Karosiene, E.; Rasmussen, M.; Blicher, T.; Lund, O.; Buus, S.; Nielsen, M. NetMHCIIpan-3.0, a Common Pan-Specific MHC Class II Prediction Method Including All Three Human MHC Class II Isotypes, HLA-DR, HLA-DP and HLA-DQ. Immunogenetics 2013, 65 (10), 711–724. https://doi.org/10.1007/s00251-013-0720-y.

23. Nielsen, M.; Lundegaard, C.; Blicher, T.; Lamberth, K.; Harndahl, M.; Justesen, S.; Røder, G.; Peters, B.; Sette, A.; Lund, O.;, et al. NetMHCpan, a Method for Quantitative Predictions of Peptide Binding to Any HLA-A and -B Locus Protein of Known Sequence. PloS One 2007, 2 (8), e796. https://doi.org/10.1371/journal.pone.0000796.

24. Abelin, J. G.; Keskin, D. B.; Sarkizova, S.; Hartigan, C. R.; Zhang, W.; Sidney, J.; Stevens, J.; Lane, W.; Zhang, G. L.; Eisenhaure, T. M.;, et al. Mass Spectrometry Profiling of HLA-Associated Peptidomes in Mono-Allelic Cells Enables More Accurate Epitope Prediction. Immunity 2017, 46 (2), 315–326. https://doi.org/10.1016/j.immuni.2017.02.007.

25. Abelin, J. G.; Harjanto, D.; Malloy, M.; Suri, P.; Colson, T.; Goulding, S. P.; Creech, A. L.; Serrano, L. R.; Nasir, G.; Nasrullah, Y.;, et al. Defining HLA-II Ligand Processing and Binding Rules with Mass Spectrometry Enhances Cancer Epitope Prediction. Immunity 2019, 51 (4), 766–779.e17. https://doi.org/10.1016/j.immuni.2019.08.012.

26. Andreatta, M.; Alvarez, B.; Nielsen, M. GibbsCluster: Unsupervised Clustering and Alignment of Peptide Sequences. Nucleic Acids Res. 2017, 45 (W1), W458–W463. https://doi.org/10.1093/nar/gkx248.

27. Bassani-Sternberg, M.; Gfeller, D. Unsupervised HLA Peptidome Deconvolution Improves Ligand Prediction Accuracy and Predicts Cooperative Effects in Peptide-HLA Interactions. J. Immunol. Baltim. Md 1950 2016, 197 (6), 2492–2499. https://doi.org/10.4049/jimmunol.1600808.

28. Alvarez, B.; Reynisson, B.; Barra, C.; Buus, S.; Ternette, N.; Connelley, T.; Andreatta, M.; Nielsen, M. NNAlign_MA; MHC Peptidome Deconvolution for Accurate MHC Binding Motif Characterization and Improved T Cell Epitope Predictions. Mol. Cell. Proteomics MCP 2019. https://doi.org/10.1074/mcp.TIR119.001658.

29. Jensen, K. K.; Andreatta, M.; Marcatili, P.; Buus, S.; Greenbaum, J. A.; Yan, Z.; Sette, A.; Peters, B.; Nielsen, M. Improved Methods for Predicting Peptide Binding Affinity to MHC Class II Molecules. Immunology 2018, 154 (3), 394–406. https://doi.org/10.1111/imm.12889.

30. Nielsen, M.; Lundegaard, C.; Worning, P.; Lauemøller, S. L.; Lamberth, K.; Buus, S.; Brunak, S.; Lund, O. Reliable Prediction of T-Cell Epitopes Using Neural Networks with Novel Sequence Representations. Protein Sci. Publ. Protein Soc. 2003, 12 (5), 1007–1017. https://doi.org/10.1110/ps.0239403.

31. Vita, R.; Mahajan, S.; Overton, J. A.; Dhanda, S. K.; Martini, S.; Cantrell, J. R.; Wheeler, D. K.; Sette, A.; Peters, B. The Immune Epitope Database (IEDB): 2018 Update. Nucleic Acids Res. 2019, 47 (D1), D339–D343. https://doi.org/10.1093/nar/gky1006.

32. Heyder, T.; Kohler, M.; Tarasova, N. K.; Haag, S.; Rutishauser, D.; Rivera, N. V.; Sandin, C.; Mia, S.; Malmström, V.; Wheelock, Å. M.;, et al. Approach for Identifying Human Leukocyte Antigen (HLA)-DR Bound Peptides from Scarce Clinical Samples. Mol. Cell. Proteomics MCP 2016, 15 (9), 3017–3029. https://doi.org/10.1074/mcp.M116.060764.

33. Bergseng, E.; Dørum, S.; Arntzen, M. Ø.; Nielsen, M.; Nygård, S.; Buus, S.; de Souza, G. A.; Sollid, L. M. Different Binding Motifs of the Celiac Disease-Associated HLA Molecules DQ2.5, DQ2.2, and DQ7.5 Revealed by Relative Quantitative Proteomics of Endogenous Peptide Repertoires. Immunogenetics 2015, 67 (2), 73–84. https://doi.org/10.1007/s00251-014-0819-9.

34. Sofron, A.; Ritz, D.; Neri, D.; Fugmann, T. High-Resolution Analysis of the Murine MHC Class II Immunopeptidome. Eur. J. Immunol. 2016, 46 (2), 319–328. https://doi.org/10.1002/eji.201545930.

35. Clement, C. C.; Becerra, A.; Yin, L.; Zolla, V.; Huang, L.; Merlin, S.; Follenzi, A.; Shaffer, S. A.; Stern, L. J.; Santambrogio, L. The Dendritic Cell Major Histocompatibility Complex II (MHC II) Peptidome Derives from a Variety of Processing Pathways and Includes Peptides with a Broad Spectrum of HLA-DM Sensitivity. J. Biol. Chem. 2016, 291 (11), 5576–5595. https://doi.org/10.1074/jbc.M115.655738.

36. Karunakaran, K. P.; Yu, H.; Jiang, X.; Chan, Q.; Goldberg, M. F.; Jenkins, M. K.; Foster, L. J.; Brunham, R. C. Identification of MHC-Bound Peptides from Dendritic Cells Infected with Salmonella Enterica Strain SL1344: Implications for a Nontyphoidal Salmonella Vaccine. J. Proteome Res. 2017, 16 (1), 298–306. https://doi.org/10.1021/acs.jproteome.6b00926.

37. Ooi, J. D.; Petersen, J.; Tan, Y. H.; Huynh, M.; Willett, Z. J.; Ramarathinam, S. H.; Eggenhuizen, P. J.; Loh, K. L.; Watson, K. A.; Gan, P. Y.;, et al. Dominant Protection from HLA-Linked Autoimmunity by Antigen-Specific Regulatory T Cells. Nature 2017, 545 (7653), 243–247. https://doi.org/10.1038/nature22329.

38. Wang, Q.; Drouin, E. E.; Yao, C.; Zhang, J.; Huang, Y.; Leon, D. R.; Steere, A. C.; Costello, C. E. Immunogenic HLA-DR-Presented Self-Peptides Identified Directly from Clinical Samples of Synovial Tissue, Synovial Fluid, or Peripheral Blood in Patients with Rheumatoid Arthritis or Lyme Arthritis. J. Proteome Res. 2017, 16 (1), 122–136. https://doi.org/10.1021/acs.jproteome.6b00386.

39. Nelde, A.; Kowalewski, D. J.; Backert, L.; Schuster, H.; Werner, J.-O.; Klein, R.; Kohlbacher, O.; Kanz, L.; Salih, H. R.; Rammensee, H.-G.;, et al. HLA Ligandome Analysis of Primary Chronic Lymphocytic Leukemia (CLL) Cells under Lenalidomide Treatment Confirms the Suitability of Lenalidomide for Combination with T-Cell-Based Immunotherapy. Oncoimmunology 2018, 7 (4), e1316438. https://doi.org/10.1080/2162402X.2017.1316438.

40. Ritz, D.; Sani, E.; Debiec, H.; Ronco, P.; Neri, D.; Fugmann, T. Membranal and Blood-Soluble HLA Class II Peptidome Analyses Using Data-Dependent and Independent Acquisition. Proteomics 2018, 18 (12), e1700246. https://doi.org/10.1002/pmic.201700246.

41. Ting, Y. T.; Petersen, J.; Ramarathinam, S. H.; Scally, S. W.; Loh, K. L.; Thomas, R.; Suri, A.; Baker, D. G.; Purcell, A. W.; Reid, H. H.;, et al. The Interplay between Citrullination and HLA-DRB1 Polymorphism in Shaping Peptide Binding Hierarchies in Rheumatoid Arthritis. J. Biol. Chem. 2018, 293 (9), 3236–3251. https://doi.org/10.1074/jbc.RA117.001013.

42. Scally, S. W.; Petersen, J.; Law, S. C.; Dudek, N. L.; Nel, H. J.; Loh, K. L.; Wijeyewickrema, L. C.; Eckle, S. B. G.; van Heemst, J.; Pike, R. N.;, et al. A Molecular Basis for the Association of the HLA-DRB1 Locus, Citrullination, and Rheumatoid Arthritis. J. Exp. Med. 2013, 210 (12), 2569–2582. https://doi.org/10.1084/jem.20131241.

43. Khodadoust, M. S.; Olsson, N.; Wagar, L. E.; Haabeth, O. A. W.; Chen, B.; Swaminathan, K.; Rawson, K.; Liu, C. L.; Steiner, D.; Lund, P.;, et al. Antigen Presentation Profiling Reveals Recognition of Lymphoma Immunoglobulin Neoantigens. Nature 2017, 543 (7647), 723–727. https://doi.org/10.1038/nature21433.

44. Álvaro-Benito, M.; Morrison, E.; Abualrous, E. T.; Kuropka, B.; Freund, C. Quantification of HLA-DM-Dependent Major Histocompatibility Complex of Class II Immunopeptidomes by the Peptide Landscape Antigenic Epitope Alignment Utility. Front. Immunol. 2018, 9, 872. https://doi.org/10.3389/fimmu.2018.00872.

45. Nanaware, P. P.; Jurewicz, M. M.; Leszyk, J. D.; Shaffer, S. A.; Stern, L. J. HLA-DO Modulates the Diversity of the MHC-II Self-Peptidome. Mol. Cell. Proteomics MCP 2019, 18 (3), 490–503. https://doi.org/10.1074/mcp.RA118.000956.

46. UniProt Consortium. UniProt: A Worldwide Hub of Protein Knowledge. Nucleic Acids Res. 2019, 47 (D1), D506–D515. https://doi.org/10.1093/nar/gky1049.

47. Nielsen, M.; Lundegaard, C.; Lund, O. Prediction of MHC Class II Binding Affinity Using SMM-Align, a Novel Stabilization Matrix Alignment Method. BMC Bioinformatics 2007, 8, 238. https://doi.org/10.1186/1471-2105-8-238.

48. Thomsen, M. C. F.; Nielsen, M. Seq2Logo: A Method for Construction and Visualization of Amino Acid Binding Motifs and Sequence Profiles Including Sequence Weighting, Pseudo Counts and Two-Sided Representation of Amino Acid Enrichment and Depletion. Nucleic Acids Res. 2012, 40 (Web Server issue), W281–287. https://doi.org/10.1093/nar/gks469.

49. Nielsen, M.; Lundegaard, C.; Blicher, T.; Peters, B.; Sette, A.; Justesen, S.; Buus, S.; Lund, O. Quantitative Predictions of Peptide Binding to Any HLA-DR Molecule of Known Sequence: NetMHCIIpan. PLoS Comput. Biol. 2008, 4 (7), e1000107. https://doi.org/10.1371/journal.pcbi.1000107.

50. Nielsen, M.; Andreatta, M. NNAlign: A Platform to Construct and Evaluate Artificial Neural Network Models of Receptor-Ligand Interactions. Nucleic Acids Res. 2017, 45 (W1), W344–W349. https://doi.org/10.1093/nar/gkx276.

51. Shao, X. M.; Bhattacharya, R.; Huang, J.; Sivakumar, I. K. A.; Tokheim, C.; Zheng, L.; Hirsch, D.; Kaminow, B.; Omdahl, A.; Bonsack, M.;, et al. High-Throughput Prediction of MHC Class I and II Neoantigens with MHCnuggets. Cancer Immunol. Res. 2019. https://doi.org/10.1158/2326-6066.CIR-19-0464.

52. Chen, B.; Khodadoust, M. S.; Olsson, N.; Wagar, L. E.; Fast, E.; Liu, C. L.; Muftuoglu, Y.; Sworder, B. J.; Diehn, M.; Levy, R.;, et al. Predicting HLA Class II Antigen Presentation through Integrated Deep Learning. Nat. Biotechnol. 2019, 37 (11), 1332–1343. https://doi.org/10.1038/s41587-019-0280-2.

53. Karosiene, E.; Lundegaard, C.; Lund, O.; Nielsen, M. NetMHCcons: A Consensus Method for the Major Histocompatibility Complex Class I Predictions. Immunogenetics 2012, 64 (3), 177–186. https://doi.org/10.1007/s00251-011-0579-8.

54. Ciudad, M. T.; Sorvillo, N.; van Alphen, F. P.; Catalán, D.; Meijer, A. B.; Voorberg, J.; Jaraquemada, D. Analysis of the HLA-DR Peptidome from Human Dendritic Cells Reveals High Affinity Repertoires and Nonconventional Pathways of Peptide Generation. J. Leukoc. Biol. 2017, 101 (1), 15–27. https://doi.org/10.1189/jlb.6HI0216-069R.

55. Choe, Y.; Leonetti, F.; Greenbaum, D. C.; Lecaille, F.; Bogyo, M.; Brömme, D.; Ellman, J. A.; Craik, C. S. Substrate Profiling of Cysteine Proteases Using a Combinatorial Peptide Library Identifies Functionally Unique Specificities. J. Biol. Chem. 2006, 281 (18), 12824–12832. https://doi.org/10.1074/jbc.M513331200.

56. Biniossek, M. L.; Nägler, D. K.; Becker-Pauly, C.; Schilling, O. Proteomic Identification of Protease Cleavage Sites Characterizes Prime and Non-Prime Specificity of Cysteine Cathepsins B, L, and S. J. Proteome Res. 2011, 10 (12), 5363–5373. https://doi.org/10.1021/pr200621z.

