## Supplementary Materials for "Improved prediction of MHC II antigen presentation through integration and motif deconvolution of mass spectrometry MHC eluted ligand data"

### Table of contents

| | | |
|---|---|---|
| S-0 | Cover Page | Title, authors and table of contents |
| S-1 | Supplementary Figure 1) | Predicted binding core motif logos deconvoluted from all MA data sets. |
| S-2 | Supplementary Figure 2) | Evidence of the benefit of extending training data scope of MA data. |
| S-3 | Supplementary Figure 3) | Quantitative analysis of MA data deconvolution consistency. |
| S-4 | Supplementary Figure 4) | Kernel density estimate plot of consistency scores, colored by loci. |
| S-5 | Supplementary Figure 5) | PPV0.9 distributions of MHC deconvolutions. |
| S-6 | Supplementary Figure 6) | MHC II logos of alleles confidently learned by the trained model. |
| S-7 | Supplementary Figure 7) | Context logos from different MHC loci: DP, DQ, DR and H2. |
| S-8 | Supplementary Figure 8) | Unsupervised deconvolution of context motifs via Gibbs clustering. |
| S-9 | Supplementary Figure 9) | F-rank results of ICS+Tetramer benchmark split up by MHC restriction. |
| S-10-19 | Supplementary Table 1) | Metadata for EL data sets. |
| S-20-27 | Supplementary Table 2) | PPV0.9 of deconvolutions. |

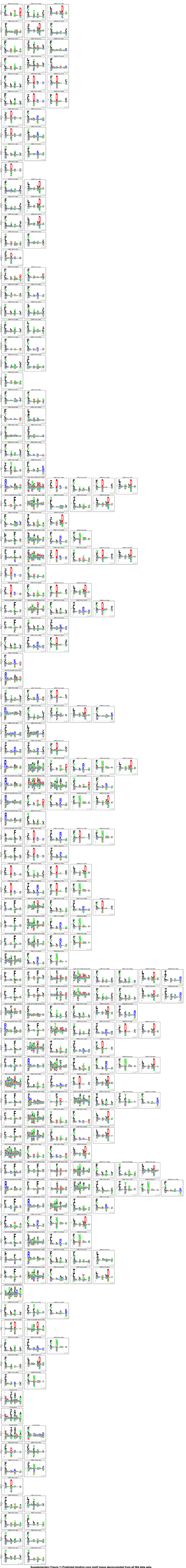

**Supplementary Figure 1) Predicted binding core motif logos deconvoluted from all MA data sets.** A single data set is shown per row and columns represent deconvolution solutions for alleles of said data set, showing the binding core logo of ligands assigned to each allele. Note that logos are generated from ligands with predicted binding >0.01 and only logos with more than 100 assigned ligands are presented.

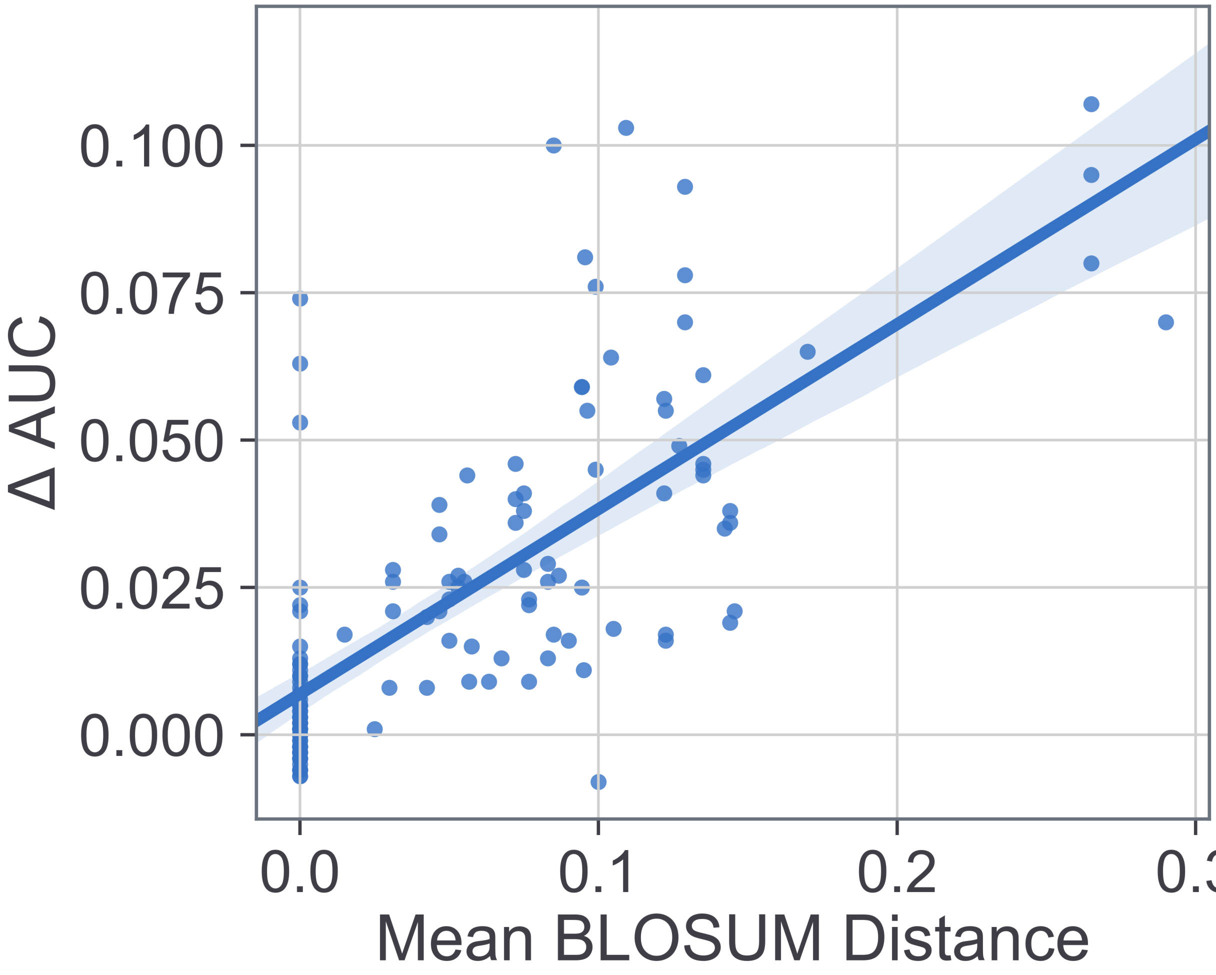


**Supplementary Figure 2) Evidence of the benefit of extending training data scope of MA data.** For cross-validation prediction performance, the MA model outperforms the SA model for datasets with alleles most distant to the SA data. This plot shows the correlation (PCC: 0.732) between AUC difference (MA-model AUC - SA model AUC) and the average allele distance of a data set from the SA data. Each point represents a single MA data set.

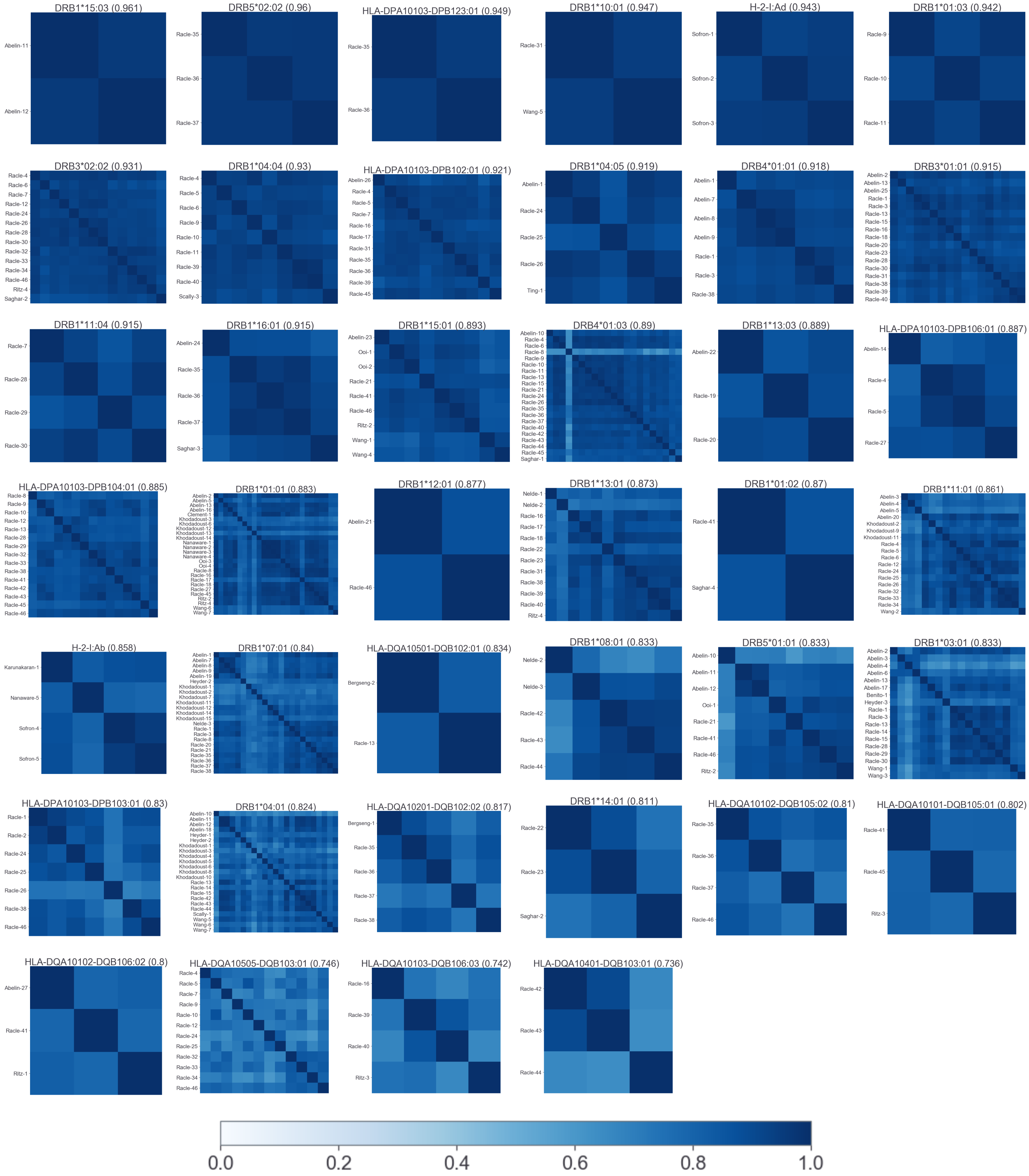


**Supplementary Figure 3) Quantitative analysis of MA data deconvolution consistency.** For each allele represented by 2 or more MA data sets, PSSM matrices for each allele were compared in terms of Pearson correlation and the resulting comparison is visualized as heatmaps. Consistency scores for each heatmap is presented.

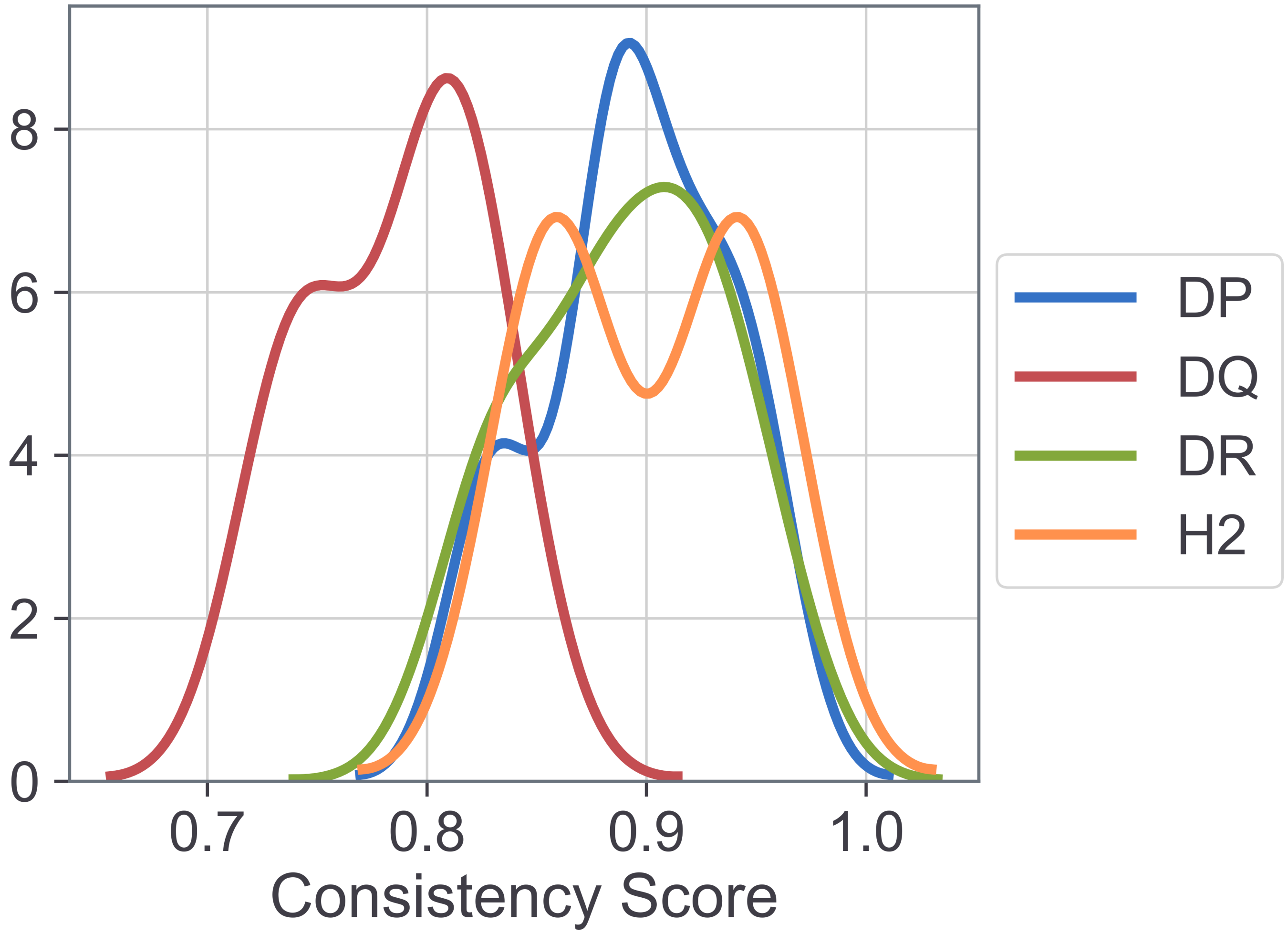


**Supplementary Figure 4) Kernel density estimate plot of consistency scores, colored by loci.** Consistency score is a quantitative estimate of agreement between logos assigned to the same MHC II molecule, across datasets.

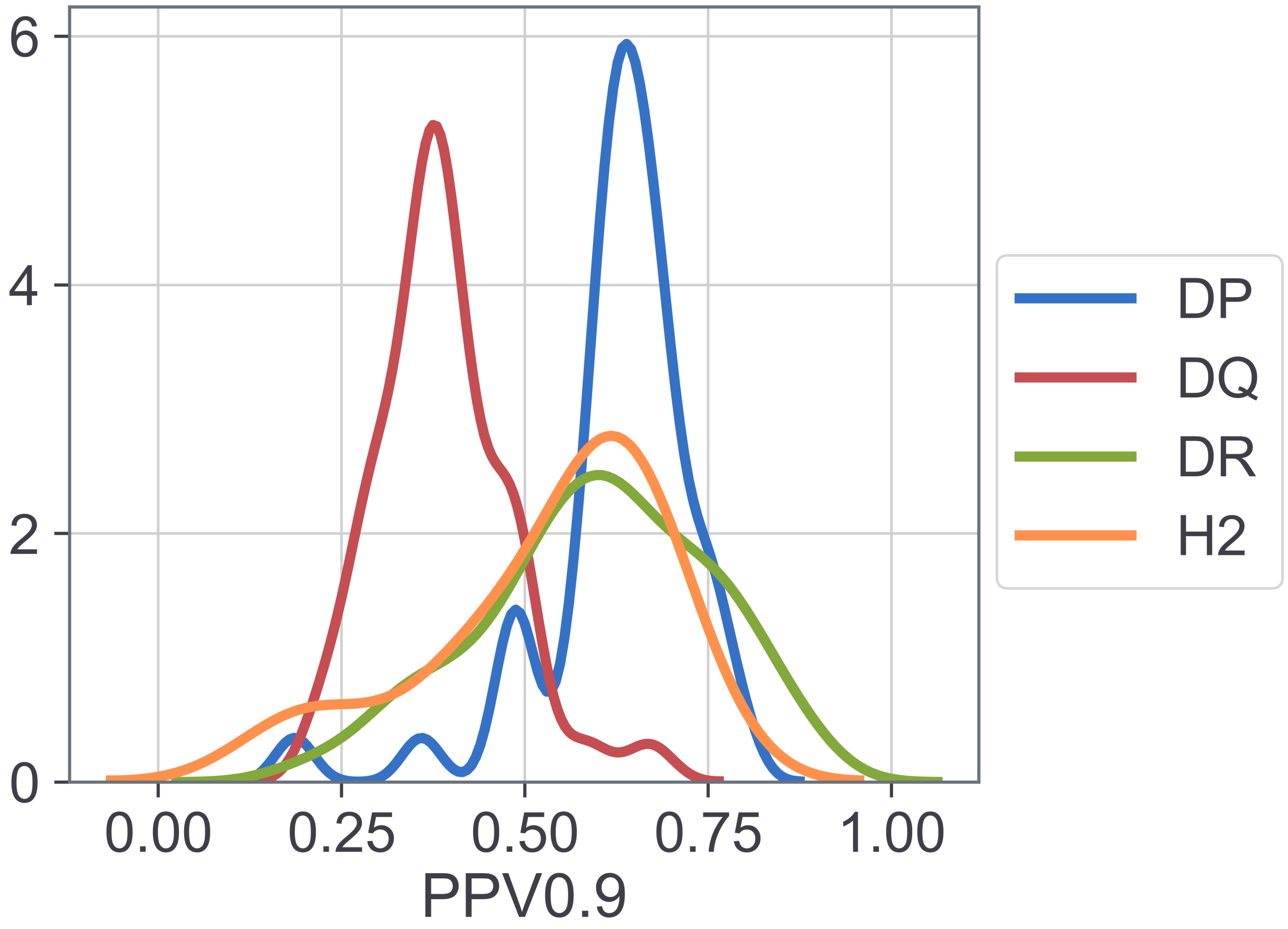


**Supplementary Figure 5) Kernel density estimate plot of the distribution of PPV0.9 for MHC deconvolutions with more than 100 ligands assigned, colored by loci.** The PPV0.9 was calculated for a 1/99 ligand/decoy datasets, for a set of decoys independent of the training set negatives.

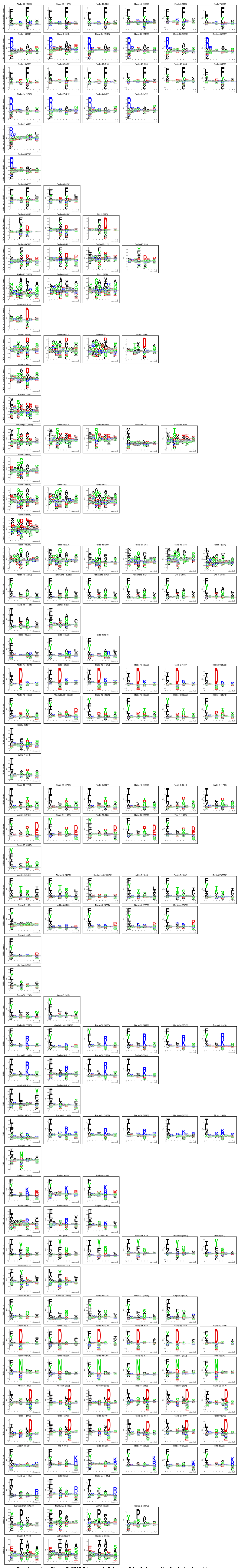

**Supplementary Figure 6) MHC II logos of alleles confidently learned by the trained model: average PPV0.9>0.3 across datasets.** Logos are compared across datasets, truncating at 6 datasets.

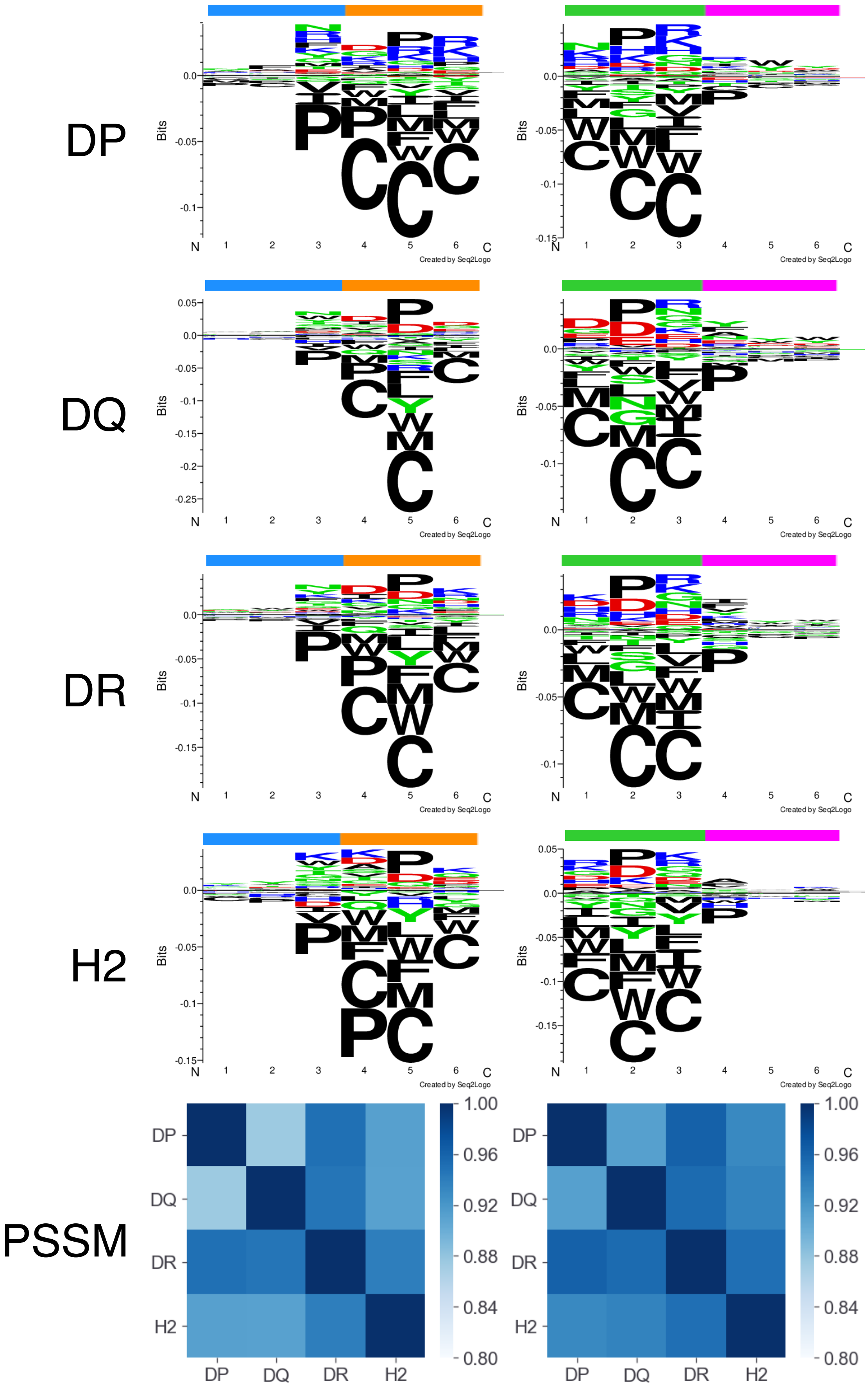


**Supplementary Figure 7) Context logos from different MHC loci: DP, DQ, DR and H2.** The figure is split down the middle, the left panel showing logos for the N-terminus and the right side showing the C-terminus. All logos share an enrichment for Proline in sites N+2 or C-2.

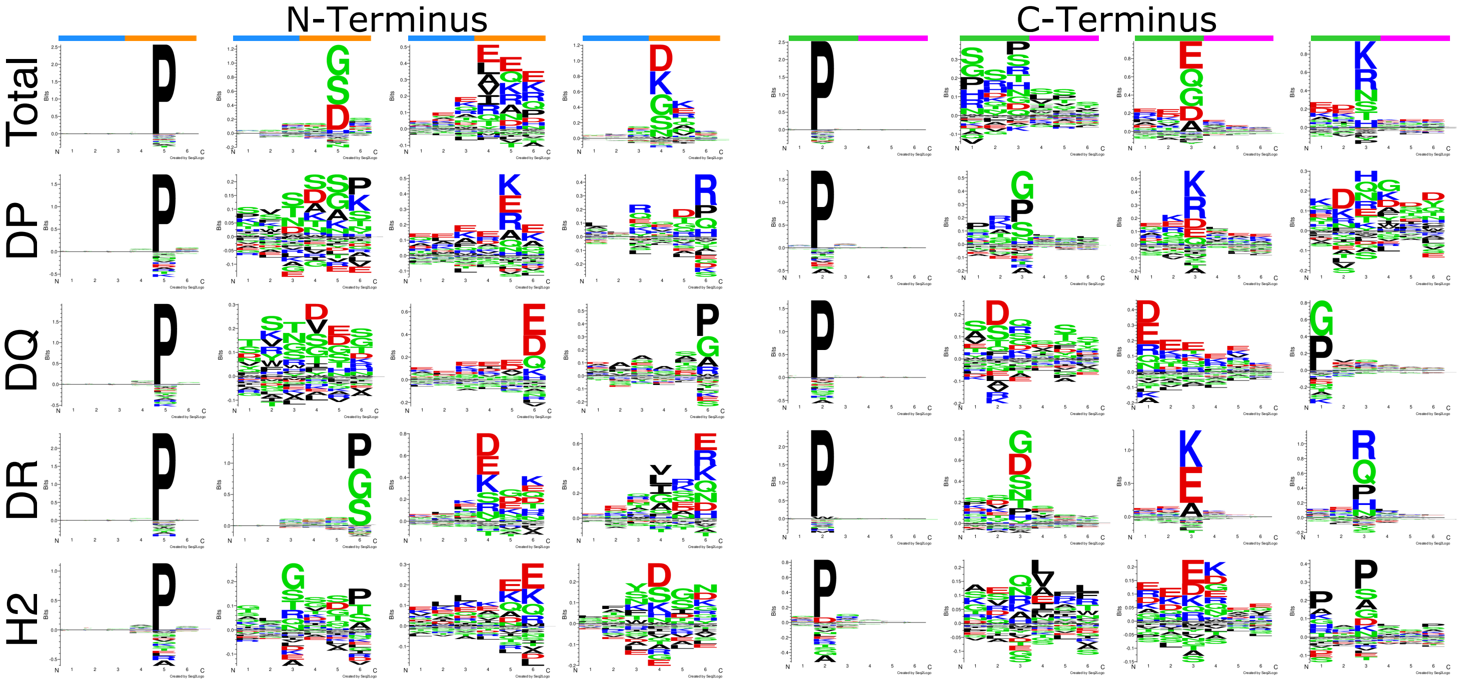


**Supplementary Figure 8) Unsupervised deconvolution of context motifs via Gibbs clustering.** Context of all loci and total set of context sequences deconvolve into similar motifs. Ordered by visual inspection.

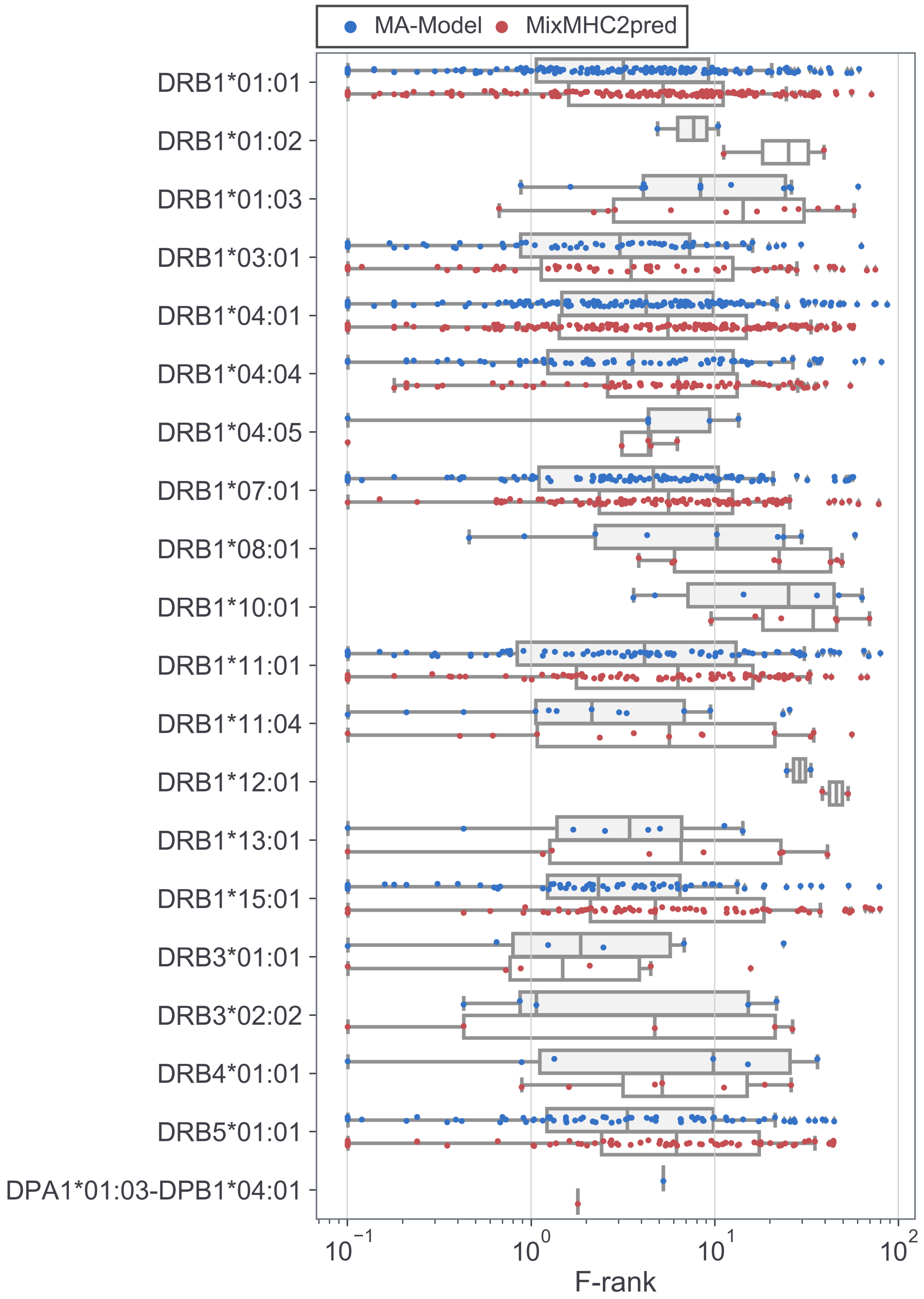


**Supplementary Figure 9) F-rank results of ICS+Tetramer benchmark split up by MHC restriction, comparing performance of the MA-Model and MixMHC2pred.** Each point represents an F-rank for a single epitope. Only epitopes of MHC molecules covered by MixMHC2pred are presented. The MA-Model has a lower median F-rank than MixMHC2pred for 16 molecules out of 20, with 1 tie (p-value<0.0045 in a binomial test).

**Supplementary Table 1) Metadata for EL data sets.**

| Cell Line ID | Dataset | Ligands | Negatives | Alleles | PMID |
|---|---|---|---|---|---|
| **HLA_DR_A375** | Abelin-1 | 3842 | 38992 | DRB1*04:05, DRB1*07:01, DRB4*01:01 | 31495665 |
| **HLA_DR_Lung** | Abelin-2 | 3364 | 32476 | DRB1*01:01, DRB1*03:01, DRB3*01:01 | 31495665 |
| **HLA_DR_PBMC_HDSC** | Abelin-3 | 339 | 3913 | DRB1*03:01, DRB1*11:01, DRB3*01:01, DRB3*02:02 | 31495665 |
| **HLA_DR_PBMC_RG1095** | Abelin-4 | 274 | 3105 | DRB1*03:01, DRB1*11:01, DRB3*01:01, DRB3*02:02 | 31495665 |
| **HLA_DR_PBMC_RG1104** | Abelin-5 | 471 | 5893 | DRB1*01:01, DRB1*11:01, DRB3*02:02 | 31495665 |
| **HLA_DR_PBMC_RG1248** | Abelin-6 | 411 | 4319 | DRB1*03:01, DRB3*01:01 | 31495665 |
| **HLA_DR_SILAC_Donor1_10minLysate** | Abelin-7 | 542 | 5849 | DRB1*07:01, DRB4*01:01 | 31495665 |
| **HLA_DR_SILAC_Donor1_5hrLysate** | Abelin-8 | 665 | 7374 | DRB1*07:01, DRB4*01:01 | 31495665 |
| **HLA_DR_SILAC_Donor1_DConly** | Abelin-9 | 1220 | 11605 | DRB1*07:01, DRB4*01:01 | 31495665 |
| **HLA_DR_SILAC_Donor1_UVovernight** | Abelin-10 | 571 | 5845 | DRB1*04:01, DRB1*15:03, DRB4*01:03, DRB5*01:01 | 31495665 |
| **HLA_DR_SILAC_Donor2_DC_UV_16hr** | Abelin-11 | 1924 | 21226 | DRB1*04:01, DRB1*15:03, DRB4*01:03, DRB5*01:01 | 31495665 |
| **HLA_DR_SILAC_Donor2_DC_UV_24hr** | Abelin-12 | 1628 | 17816 | DRB1*04:01, DRB1*15:03, DRB4*01:03, DRB5*01:01 | 31495665 |
| **HLA_DR_Spleen** | Abelin-13 | 2412 | 21592 | DRB1*01:01, DRB1*03:01, DRB3*01:01 | 31495665 |
| **MAPTAC_DPB1_0601_DPA1_0103_dmPos** | Abelin-14 | 1795 | 14846 | DPA1*01:03-DPB1*06:01 | 31495665 |
| **MAPTAC_DQB1_0604_DQA1_0102_dmPos** | Abelin-15 | 981 | 7739 | DQA1*01:02-DQB1*06:04 | 31495665 |
| **MAPTAC_DRB1_0101** | Abelin-16 | 6298 | 49435 | DRB1*01:01 | 31495665 |
| **MAPTAC_DRB1_0301** | Abelin-17 | 2118 | 19118 | DRB1*03:01 | 31495665 |
| **MAPTAC_DRB1_0401** | Abelin-18 | 1960 | 17950 | DRB1*04:01 | 31495665 |
| **MAPTAC_DRB1_0701** | Abelin-19 | 4297 | 34365 | DRB1*07:01 | 31495665 |
| **MAPTAC_DRB1_1101** | Abelin-20 | 7588 | 59777 | DRB1*11:01 | 31495665 |
| **MAPTAC_DRB1_1201_dmPos** | Abelin-21 | 854 | 5981 | DRB1*12:01 | 31495665 |
| **MAPTAC_DRB1_1303** | Abelin-22 | 2677 | 22807 | DRB1*13:03 | 31495665 |
| **MAPTAC_DRB1_1501** | Abelin-23 | 2598 | 18841 | DRB1*15:01 | 31495665 |
| **MAPTAC_DRB1_1601** | Abelin-24 | 919 | 7286 | DRB1*16:01 | 31495665 |
| **MAPTAC_DRB3_0101_dmPos** | Abelin-25 | 589 | 4947 | DRB3*01:01 | 31495665 |
| **MAPTAC_HLA_DPB10201_DPA10103** | Abelin-26 | 2164 | 18667 | DPA1*01:03-DPB1*02:01 | 31495665 |

| | | | | | |
|---|---|---|---|---|---|
| **MAPTAC_HLA_DQB10 602_DQA10102** | Abelin-27 | 2950 | 29693 | DQA1*01:02-DQB1*06:02 | 31495665 |
| **T2_DR3** | Benito-1 | 1285 | 10573 | DRB1*03:01 | 29774024 |
| **DQ2_2** | Bergseng-1 | 5823 | 55729 | DQA1*02:01-DQB1*02:02 | 25502872 |
| **DQ2_5** | Bergseng-2 | 3052 | 28610 | DQA1*05:01-DQB1*02:01 | 25502872 |
| **DQ7_5** | Bergseng-3 | 6149 | 55953 | DQA1*05:01-DQB1*03:01 | 25502872 |
| **DRB1_0101** | Clement-1 | 2963 | 30362 | DRB1*01:01 | 26740625 |
| **Bcell** | Heyder-1 | 388 | 3327 | DRB1*04:01 | 27452731 |
| **PAT1** | Heyder-2 | 670 | 7738 | DRB1*04:01, DRB1*07:01 | 27452731 |
| **PAT7** | Heyder-3 | 287 | 2790 | DRB1*03:01, DRB1*15:01 | 27452731 |
| **H_2_IAb** | Karunakaran-1 | 1544 | 13180 | H-2-IAb | 27802388 |
| **Jeko** | Khodadoust-1 | 4332 | 38277 | DRB1*04:01, DRB1*07:01 | 28329770 |
| **L128** | Khodadoust-2 | 4731 | 44479 | DRB1*07:01, DRB1*11:01 | 28329770 |
| **MCL001** | Khodadoust-3 | 1925 | 19028 | DRB1*01:01, DRB1*04:01 | 28329770 |
| **MCL005** | Khodadoust-4 | 401 | 4545 | DRB1*04:01, DRB1*11:01 | 28329770 |
| **MCL008** | Khodadoust-5 | 672 | 6882 | DRB1*04:01 | 28329770 |
| **MCL022** | Khodadoust-6 | 920 | 10211 | DRB1*01:01, DRB1*04:01 | 28329770 |
| **MCL030** | Khodadoust-7 | 1095 | 10482 | DRB1*07:01 | 28329770 |
| **MCL034** | Khodadoust-8 | 628 | 8992 | DRB1*04:01 | 28329770 |
| **MCL037** | Khodadoust-9 | 1218 | 12054 | DRB1*11:01 | 28329770 |
| **MCL041** | Khodadoust-10 | 814 | 8321 | DRB1*04:01 | 28329770 |
| **MCL043** | Khodadoust-11 | 1173 | 11651 | DRB1*07:01, DRB1*11:01 | 28329770 |
| **MCL049** | Khodadoust-12 | 453 | 5717 | DRB1*01:01, DRB1*07:01 | 28329770 |
| **MCL052** | Khodadoust-13 | 986 | 10349 | DRB1*01:01 | 28329770 |
| **MCLX001** | Khodadoust-14 | 784 | 8231 | DRB1*01:01, DRB1*07:01 | 28329770 |
| **MCLX002** | Khodadoust-15 | 373 | 3689 | DRB1*07:01 | 28329770 |
| **DO_KO_1** | Nanaware-1 | 3596 | 34637 | DRB1*01:01 | 30573663 |
| **DO_KO_2** | Nanaware-2 | 2455 | 25008 | DRB1*01:01 | 30573663 |
| **WT_1** | Nanaware-3 | 4418 | 45301 | DRB1*01:01 | 30573663 |
| **WT_2** | Nanaware-4 | 3207 | 33328 | DRB1*01:01 | 30573663 |
| **WT_H2** | Nanaware-5 | 481 | 4140 | H-2-IAb | 30573663 |
| **UPN1** | Nelde-1 | 3079 | 29736 | DRB1*08:03, DRB1*13:01 | 29632711 |
| **UPN2** | Nelde-2 | 367 | 3555 | DRB1*08:01, DRB1*13:01 | 29632711 |
| **UPN3** | Nelde-3 | 3216 | 29552 | DRB1*07:01, DRB1*08:01 | 29632711 |
| **DR15_human** | Ooi-1 | 2425 | 22358 | DRB1*15:01, DRB5*01:01 | 28467828 |
| **DR15_mouse** | Ooi-2 | 3327 | 36605 | DRB1*15:01 | 28467828 |
| **DR1_human** | Ooi-3 | 4254 | 37421 | DRB1*01:01 | 28467828 |
| **DR1_mouse** | Ooi-4 | 3986 | 44566 | DRB1*01:01 | 28467828 |
| **3808_HMC** | Racle-1 | 5262 | 56629 | DPA1*01:03-DPB1*03:01, DPA1*01:03-DPB1*11:01, DPA1*02:01-DPB1*03:01, DPA1*02:01-DPB1*11:01, DQA1*02:01-DQB1*02:01, DQA1*02:01-DQB1*02:02, DQA1*05:01-DQB1*02:01, | 31611696 |

| | | | | | |
|---|---|---|---|---|---|
| | | | | DQA1*05:01-DQB1*02:02, DRB1*03:01, DRB1*07:01, DRB3*01:01, DRB4*01:01 | |
| **3808_HMC_DQP** | Racle-2 | 1002 | 11427 | DPA1*01:03-DPB1*03:01, DPA1*01:03-DPB1*11:01, DPA1*02:01-DPB1*03:01, DPA1*02:01-DPB1*11:01, DQA1*02:01-DQB1*02:01, DQA1*02:01-DQB1*02:02, DQA1*05:01-DQB1*02:01, DQA1*05:01-DQB1*02:02, DRB1*03:01, DRB1*07:01, DRB3*01:01, DRB4*01:01 | 31611696 |
| **3808_HMC_DR** | Racle-3 | 4077 | 44520 | DPA1*01:03-DPB1*03:01, DPA1*01:03-DPB1*11:01, DPA1*02:01-DPB1*03:01, DPA1*02:01-DPB1*11:01, DQA1*02:01-DQB1*02:01, DQA1*02:01-DQB1*02:02, DQA1*05:01-DQB1*02:01, DQA1*05:01-DQB1*02:02, DRB1*03:01, DRB1*07:01, DRB3*01:01, DRB4*01:01 | 31611696 |
| **3830NJF** | Racle-4 | 8044 | 81407 | DPA1*01:03-DPB1*02:01, DPA1*01:03-DPB1*06:01, DQA1*03:01-DQB1*03:01, DQA1*03:01-DQB1*03:02, DQA1*05:05-DQB1*03:01, DQA1*05:05-DQB1*03:02, DRB1*04:04, DRB1*11:01, DRB3*02:02, DRB4*01:03 | 31611696 |
| **3830_NJF_DQP** | Racle-5 | 3503 | 36877 | DPA1*01:03-DPB1*02:01, DPA1*01:03-DPB1*06:01, DQA1*03:01-DQB1*03:01, DQA1*03:01-DQB1*03:02, DQA1*05:05-DQB1*03:01, DQA1*05:05-DQB1*03:02, DRB1*04:04, DRB1*11:01, DRB3*02:02, DRB4*01:03 | 31611696 |
| **3830_NJF_DR** | Racle-6 | 2781 | 29281 | DPA1*01:03-DPB1*02:01, DPA1*01:03-DPB1*06:01, DQA1*03:01-DQB1*03:01, DQA1*03:01-DQB1*03:02, DQA1*05:05-DQB1*03:01, DQA1*05:05-DQB1*03:02, DRB1*04:04, DRB1*11:01, DRB3*02:02, DRB4*01:03 | 31611696 |
| **3849BR** | Racle-7 | 7292 | 74963 | DPA1*01:03-DPB1*02:01, DPA1*01:03-DPB1*04:01, | 31611696 |

| | | | | | |
|---|---|---|---|---|---|
| | | | | DQA1*05:05-DQB1*03:01, DRB1*11:04, DRB3*02:02 | |
| **3865DM** | Racle-8 | 4510 | 44847 | DPA1*01:03-DPB1*04:01, DPA1*01:03-DPB1*20:01, DQA1*01:01-DQB1*03:03, DQA1*01:01-DQB1*05:01, DQA1*02:01-DQB1*03:03, DQA1*02:01-DQB1*05:01, DRB1*01:01, DRB1*07:01, DRB4*01:03 | 31611696 |
| **3869_GA** | Racle-9 | 5365 | 55688 | DPA1*01:03-DPB1*04:01, DPA1*01:03-DPB1*12:601, DQA1*03:01-DQB1*03:01, DQA1*03:01-DQB1*03:02, DQA1*05:05-DQB1*03:01, DQA1*05:05-DQB1*03:02, DRB1*01:03, DRB1*04:04, DRB4*01:03 | 31611696 |
| **3869_GA_DQP** | Racle-10 | 2203 | 23749 | DPA1*01:03-DPB1*04:01, DPA1*01:03-DPB1*12:601, DQA1*03:01-DQB1*03:01, DQA1*03:01-DQB1*03:02, DQA1*05:05-DQB1*03:01, DQA1*05:05-DQB1*03:02, DRB1*01:03, DRB1*04:04, DRB4*01:03 | 31611696 |
| **3869_GA_DR** | Racle-11 | 3218 | 35036 | DPA1*01:03-DPB1*04:01, DPA1*01:03-DPB1*12:601, DQA1*03:01-DQB1*03:01, DQA1*03:01-DQB1*03:02, DQA1*05:05-DQB1*03:01, DQA1*05:05-DQB1*03:02, DRB1*01:03, DRB1*04:04, DRB4*01:03 | 31611696 |
| **3911_ME** | Racle-12 | 2135 | 21637 | DPA1*01:03-DPB1*04:01, DQA1*05:05-DQB1*03:01, DRB1*11:01, DRB3*02:02 | 31611696 |
| **3912BAM** | Racle-13 | 4621 | 48212 | DPA1*01:03-DPB1*04:01, DQA1*03:01-DQB1*02:01, DQA1*03:01-DQB1*03:02, DQA1*05:01-DQB1*02:01, DQA1*05:01-DQB1*03:02, DRB1*03:01, DRB1*04:01, DRB3*01:01, DRB4*01:03 | 31611696 |
| **3912_BAM_DQP** | Racle-14 | 552 | 6387 | DPA1*01:03-DPB1*04:01, DQA1*03:01-DQB1*02:01, DQA1*03:01-DQB1*03:02, DQA1*05:01-DQB1*02:01, DQA1*05:01-DQB1*03:02, | 31611696 |

| | | | | | |
|---|---|---|---|---|---|
| | | | | DRB1*03:01, DRB1*04:01, DRB3*01:01, DRB4*01:03 | |
| **3912_BAM_DR** | Racle-15 | 5768 | 56844 | DPA1*01:03-DPB1*04:01, DQA1*03:01-DQB1*02:01, DQA1*03:01-DQB1*03:02, DQA1*05:01-DQB1*02:01, DQA1*05:01-DQB1*03:02, DRB1*03:01, DRB1*04:01, DRB3*01:01, DRB4*01:03 | 31611696 |
| **3947_GA** | Racle-16 | 4545 | 47050 | DPA1*01:03-DPB1*02:01, DPA1*01:03-DPB1*04:02, DQA1*01:01-DQB1*05:01, DQA1*01:01-DQB1*06:03, DQA1*01:03-DQB1*05:01, DQA1*01:03-DQB1*06:03, DRB1*01:01, DRB1*13:01, DRB3*01:01 | 31611696 |
| **3947_GA_DQP** | Racle-17 | 791 | 9041 | DPA1*01:03-DPB1*02:01, DPA1*01:03-DPB1*04:02, DQA1*01:01-DQB1*05:01, DQA1*01:01-DQB1*06:03, DQA1*01:03-DQB1*05:01, DQA1*01:03-DQB1*06:03, DRB1*01:01, DRB1*13:01, DRB3*01:01 | 31611696 |
| **3947_GA_DR** | Racle-18 | 3241 | 33689 | DPA1*01:03-DPB1*02:01, DPA1*01:03-DPB1*04:02, DQA1*01:01-DQB1*05:01, DQA1*01:01-DQB1*06:03, DQA1*01:03-DQB1*05:01, DQA1*01:03-DQB1*06:03, DRB1*01:01, DRB1*13:01, DRB3*01:01 | 31611696 |
| **3971_ORA** | Racle-19 | 357 | 4048 | DPA1*01:03-DPB1*04:01, DPA1*02:02-DPB1*04:01, DQA1*02:01-DQB1*02:02, DQA1*02:01-DQB1*03:01, DQA1*05:05-DQB1*02:02, DQA1*05:05-DQB1*03:01, DRB1*07:01, DRB1*13:03, DRB3*01:01, DRB4*01:01 | 31611696 |
| **3971_ORA_DR** | Racle-20 | 1391 | 13588 | DPA1*01:03-DPB1*04:01, DPA1*02:02-DPB1*04:01, DQA1*02:01-DQB1*02:02, DQA1*02:01-DQB1*03:01, DQA1*05:05-DQB1*02:02, DQA1*05:05-DQB1*03:01, DRB1*07:01, DRB1*13:03, DRB3*01:01, DRB4*01:01 | 31611696 |

| | | | | | |
|---|---|---|---|---|---|
| **3993** | Racle-21 | 1647 | 15029 | DPA1*01:03-DPB1*04:01, DPA1*01:03-DPB1*17:01, DPA1*02:01-DPB1*04:01, DPA1*02:01-DPB1*17:01, DQA1*01:02-DQB1*02:02, DQA1*01:02-DQB1*06:02, DQA1*02:01-DQB1*02:02, DQA1*02:01-DQB1*06:02, DRB1*07:01, DRB1*15:01, DRB4*01:03, DRB5*01:01 | 31611696 |
| **4001** | Racle-22 | 322 | 3373 | DPA1*01:03-DPB1*04:01, DPA1*01:03-DPB1*04:02, DQA1*01:03-DQB1*05:03, DQA1*01:03-DQB1*06:03, DQA1*01:04-DQB1*05:03, DQA1*01:04-DQB1*06:03, DRB1*13:01, DRB1*14:01, DRB3*01:01, DRB3*02:02 | 31611696 |
| **4001_DR** | Racle-23 | 1362 | 13450 | DPA1*01:03-DPB1*04:01, DPA1*01:03-DPB1*04:02, DQA1*01:03-DQB1*05:03, DQA1*01:03-DQB1*06:03, DQA1*01:04-DQB1*05:03, DQA1*01:04-DQB1*06:03, DRB1*13:01, DRB1*14:01, DRB3*01:01, DRB3*02:02 | 31611696 |
| **4021** | Racle-24 | 5857 | 60776 | DPA1*01:03-DPB1*03:01, DPA1*01:03-DPB1*10:401, DQA1*03:03-DQB1*02:02, DQA1*03:03-DQB1*03:01, DQA1*05:05-DQB1*02:02, DQA1*05:05-DQB1*03:01, DRB1*04:05, DRB1*11:01, DRB3*02:02, DRB4*01:03 | 31611696 |
| **4021_DQP** | Racle-25 | 3802 | 38414 | DPA1*01:03-DPB1*03:01, DPA1*01:03-DPB1*10:401, DQA1*03:03-DQB1*02:02, DQA1*03:03-DQB1*03:01, DQA1*05:05-DQB1*02:02, DQA1*05:05-DQB1*03:01, DRB1*04:05, DRB1*11:01, DRB3*02:02, DRB4*01:03 | 31611696 |
| **4021_DR** | Racle-26 | 4620 | 48129 | DPA1*01:03-DPB1*03:01, DPA1*01:03-DPB1*10:401, DQA1*03:03-DQB1*02:02, DQA1*03:03-DQB1*03:01, DQA1*05:05-DQB1*02:02, DQA1*05:05-DQB1*03:01, DRB1*04:05, DRB1*11:01, DRB3*02:02, DRB4*01:03 | 31611696 |

| | | | | | |
|---|---|---|---|---|---|
| **4037_DC** | Racle-27 | 2668 | 27571 | DPA1*01:03-DPB1*04:01, DPA1*01:03-DPB1*06:01, DQA1*01:01-DQB1*05:01, DRB1*01:01 | 31611696 |
| **4052_BA** | Racle-28 | 3837 | 38993 | DPA1*01:03-DPB1*04:01, DQA1*05:01-DQB1*02:01, DQA1*05:01-DQB1*03:01, DQA1*05:05-DQB1*02:01, DQA1*05:05-DQB1*03:01, DRB1*03:01, DRB1*11:04, DRB3*01:01, DRB3*02:02 | 31611696 |
| **4052_BA_DQP** | Racle-29 | 714 | 7329 | DPA1*01:03-DPB1*04:01, DQA1*05:01-DQB1*02:01, DQA1*05:01-DQB1*03:01, DQA1*05:05-DQB1*02:01, DQA1*05:05-DQB1*03:01, DRB1*03:01, DRB1*11:04, DRB3*01:01, DRB3*02:02 | 31611696 |
| **4052_BA_DR** | Racle-30 | 5367 | 54593 | DPA1*01:03-DPB1*04:01, DQA1*05:01-DQB1*02:01, DQA1*05:01-DQB1*03:01, DQA1*05:05-DQB1*02:01, DQA1*05:05-DQB1*03:01, DRB1*03:01, DRB1*11:04, DRB3*01:01, DRB3*02:02 | 31611696 |
| **BP455** | Racle-31 | 5323 | 48707 | DPA1*01:03-DPB1*02:01, DQA1*01:05-DQB1*05:01, DQA1*01:05-DQB1*06:03, DQA1*01:10-DQB1*05:01, DQA1*01:10-DQB1*06:03, DRB1*10:01, DRB1*13:01, DRB3*01:01 | 31611696 |
| **CD165** | Racle-32 | 8210 | 74439 | DPA1*01:03-DPB1*04:01, DPA1*01:03-DPB1*04:02, DQA1*05:05-DQB1*03:01, DRB1*11:01, DRB3*02:02 | 31611696 |
| **CD165_DQP** | Racle-33 | 6309 | 61892 | DPA1*01:03-DPB1*04:01, DPA1*01:03-DPB1*04:02, DQA1*05:05-DQB1*03:01, DRB1*11:01, DRB3*02:02 | 31611696 |
| **CD165_DR** | Racle-34 | 7905 | 72587 | DPA1*01:03-DPB1*04:01, DPA1*01:03-DPB1*04:02, DQA1*05:05-DQB1*03:01, DRB1*11:01, DRB3*02:02 | 31611696 |
| **CM647** | Racle-35 | 8001 | 77135 | DPA1*01:03-DPB1*02:01, DPA1*01:03-DPB1*23:01, DQA1*01:02-DQB1*02:02, DQA1*01:02-DQB1*05:02, DQA1*02:01-DQB1*02:02, DQA1*02:01-DQB1*05:02, | 31611696 |

| | | | | | |
|---|---|---|---|---|---|
| | | | | DRB1*07:01, DRB1*16:01, DRB4*01:03, DRB5*02:02 | |
| **CM647_DQP** | Racle-36 | 4310 | 40075 | DPA1*01:03-DPB1*02:01, DPA1*01:03-DPB1*23:01, DQA1*01:02-DQB1*02:02, DQA1*01:02-DQB1*05:02, DQA1*02:01-DQB1*02:02, DQA1*02:01-DQB1*05:02, DRB1*07:01, DRB1*16:01, DRB4*01:03, DRB5*02:02 | 31611696 |
| **CM647_DR** | Racle-37 | 6112 | 57213 | DPA1*01:03-DPB1*02:01, DPA1*01:03-DPB1*23:01, DQA1*01:02-DQB1*02:02, DQA1*01:02-DQB1*05:02, DQA1*02:01-DQB1*02:02, DQA1*02:01-DQB1*05:02, DRB1*07:01, DRB1*16:01, DRB4*01:03, DRB5*02:02 | 31611696 |
| **GD149** | Racle-38 | 6022 | 60358 | DPA1*01:03-DPB1*03:01, DPA1*01:03-DPB1*04:01, DPA1*02:01-DPB1*03:01, DPA1*02:01-DPB1*04:01, DQA1*01:10-DQB1*02:02, DQA1*01:10-DQB1*06:03, DQA1*02:01-DQB1*02:02, DQA1*02:01-DQB1*06:03, DRB1*07:01, DRB1*13:01, DRB3*01:01, DRB4*01:01 | 31611696 |
| **JY** | Racle-39 | 4850 | 50146 | DPA1*01:03-DPB1*02:01, DPA1*01:03-DPB1*04:01, DQA1*01:03-DQB1*03:02, DQA1*01:03-DQB1*06:03, DQA1*03:01-DQB1*03:02, DQA1*03:01-DQB1*06:03, DRB1*04:04, DRB1*13:01, DRB3*01:01, DRB4*01:03 | 31611696 |
| **JY_DR** | Racle-40 | 4240 | 46727 | DPA1*01:03-DPB1*02:01, DPA1*01:03-DPB1*04:01, DQA1*01:03-DQB1*03:02, DQA1*01:03-DQB1*06:03, DQA1*03:01-DQB1*03:02, DQA1*03:01-DQB1*06:03, DRB1*04:04, DRB1*13:01, DRB3*01:01, DRB4*01:03 | 31611696 |
| **PD42** | Racle-41 | 8906 | 86073 | DPA1*01:03-DPB1*04:01, DPA1*01:03-DPB1*05:01, DPA1*02:02-DPB1*04:01, DPA1*02:02-DPB1*05:01, DQA1*01:01-DQB1*05:01, DQA1*01:01-DQB1*06:02, | 31611696 |

| | | | | | |
|---|---|---|---|---|---|
| | | | | DQA1*01:02-DQB1*05:01, DQA1*01:02-DQB1*06:02, DRB1*01:02, DRB1*15:01, DRB5*01:01 | |
| **RA957** | Racle-42 | 10224 | 90451 | DPA1*01:03-DPB1*04:01, DPA1*01:03-DPB1*04:02, DQA1*03:03-DQB1*03:01, DQA1*03:03-DQB1*04:02, DQA1*04:01-DQB1*03:01, DQA1*04:01-DQB1*04:02, DRB1*04:01, DRB1*08:01, DRB4*01:03 | 31611696 |
| **RA957_DQP** | Racle-43 | 4807 | 45615 | DPA1*01:03-DPB1*04:01, DPA1*01:03-DPB1*04:02, DQA1*03:03-DQB1*03:01, DQA1*03:03-DQB1*04:02, DQA1*04:01-DQB1*03:01, DQA1*04:01-DQB1*04:02, DRB1*04:01, DRB1*08:01, DRB4*01:03 | 31611696 |
| **RA957_DR** | Racle-44 | 4749 | 44390 | DPA1*01:03-DPB1*04:01, DPA1*01:03-DPB1*04:02, DQA1*03:03-DQB1*03:01, DQA1*03:03-DQB1*04:02, DQA1*04:01-DQB1*03:01, DQA1*04:01-DQB1*04:02, DRB1*04:01, DRB1*08:01, DRB4*01:03 | 31611696 |
| **TIL1** | Racle-45 | 6778 | 70572 | DPA1*01:03-DPB1*02:01, DPA1*01:03-DPB1*04:01, DQA1*01:01-DQB1*03:01, DQA1*01:01-DQB1*05:01, DQA1*03:03-DQB1*03:01, DQA1*03:03-DQB1*05:01, DRB1*01:01, DRB1*04:08, DRB4*01:03 | 31611696 |
| **TIL3** | Racle-46 | 8654 | 77798 | DPA1*01:03-DPB1*03:01, DPA1*01:03-DPB1*04:01, DQA1*01:02-DQB1*03:01, DQA1*01:02-DQB1*05:02, DQA1*05:05-DQB1*03:01, DQA1*05:05-DQB1*05:02, DRB1*12:01, DRB1*15:01, DRB3*02:02, DRB5*01:01 | 31611696 |
| **DOHH2_DQ** | Ritz-1 | 414 | 4183 | DQA1*01:01-DQB1*05:01, DQA1*01:02-DQB1*06:02 | 29314611 |
| **DOHH2_DR** | Ritz-2 | 2793 | 29010 | DRB1*01:01, DRB1*15:01, DRB5*01:01 | 29314611 |
| **Maver_1_DQ** | Ritz-3 | 2012 | 20202 | DQA1*01:01-DQB1*05:01, DQA1*01:03-DQB1*06:03 | 29314611 |

| | | | | | |
|---|---|---|---|---|---|
| **Maver_1_DR** | Ritz-4 | 5557 | 52401 | DRB1*01:01, DRB1*13:01, DRB3*02:02 | 29314611 |
| **peptide_0901** | Saghar-1 | 955 | 10207 | DRB1*09:01, DRB4*01:03 | In House Data |
| **peptide_1401** | Saghar-2 | 2229 | 21413 | DRB1*14:01, DRB3*02:02 | In House Data |
| **peptide_1601** | Saghar-3 | 1385 | 12507 | DRB1*16:01 | In House Data |
| **peptide_HeLa** | Saghar-4 | 568 | 5397 | DRB1*01:02 | In House Data |
| **DRB1_0401** | Scally-1 | 771 | 6524 | DRB1*04:01 | 24190431 |
| **DRB1_0402** | Scally-2 | 1083 | 8323 | DRB1*04:02 | 24190431 |
| **DRB1_0404** | Scally-3 | 1809 | 14533 | DRB1*04:04 | 24190431 |
| **A20** | Sofron-1 | 1214 | 12638 | H-2-IAd, H-2-IEd | 26495903 |
| **Balb_c_1** | Sofron-2 | 888 | 9536 | H-2-IAd, H-2-IEd | 26495903 |
| **Balb_c_2** | Sofron-3 | 2727 | 28747 | H-2-IAd, H-2-IEd | 26495903 |
| **C57BL_6_1** | Sofron-4 | 734 | 8137 | H-2-IAb | 26495903 |
| **C57BL_6_2** | Sofron-5 | 2493 | 26578 | H-2-IAb | 26495903 |
| **T2_DRB1_0405** | Ting-1 | 1665 | 13269 | DRB1*04:05 | 29317506 |
| **LA3** | Wang-1 | 408 | 4139 | DRB1*03:01, DRB1*15:01 | 27726376 |
| **LA4** | Wang-2 | 257 | 2832 | DRB1*11:01, DRB1*13:02 | 27726376 |
| **LA5** | Wang-3 | 327 | 3283 | DRB1*03:01, DRB1*03:05 | 27726376 |
| **LA7** | Wang-4 | 681 | 7331 | DRB1*04:03, DRB1*15:01 | 27726376 |
| **LA8** | Wang-5 | 1771 | 17446 | DRB1*04:01, DRB1*10:01 | 27726376 |
| **RA2** | Wang-6 | 288 | 2921 | DRB1*01:01, DRB1*04:01 | 27726376 |
| **RA3** | Wang-7 | 335 | 3687 | DRB1*01:01, DRB1*04:01 | 27726376 |

**Supplementary Table 2) PPV0.9 of deconvolutions. Only results for MHC deconvolutions with more than 100 ligands assigned are shown. Ligand/decoy ratio is 1/99 for a set of decoys independent of the training data.**

| Dataset | MHC | PPV0.9 | Ligands |
|---|---|---|---|
| **Racle-13** | DRB3*01:01 | 0.899 | 165 |
| **Racle-21** | DRB4*01:03 | 0.89 | 213 |
| **Racle-28** | DRB3*01:01 | 0.883 | 181 |
| **Racle-23** | DRB3*01:01 | 0.879 | 111 |
| **Racle-30** | DRB3*01:01 | 0.875 | 373 |
| **Racle-20** | DRB3*01:01 | 0.871 | 138 |
| **Racle-15** | DRB3*01:01 | 0.871 | 371 |
| **Racle-1** | DRB3*01:01 | 0.868 | 228 |
| **Racle-24** | DRB4*01:03 | 0.864 | 245 |
| **Racle-3** | DRB3*01:01 | 0.853 | 280 |
| **Racle-18** | DRB3*01:01 | 0.845 | 302 |
| **Abelin-2** | DRB3*01:01 | 0.844 | 300 |
| **Racle-10** | DRB4*01:03 | 0.838 | 165 |
| **Racle-17** | DRB1*01:01 | 0.833 | 107 |
| **Racle-14** | DRB1*03:01 | 0.822 | 225 |
| **Abelin-13** | DRB3*01:01 | 0.815 | 180 |
| **Racle-38** | DRB3*01:01 | 0.81 | 468 |
| **Racle-43** | DRB4*01:03 | 0.809 | 123 |
| **Racle-26** | DRB4*01:03 | 0.807 | 404 |
| **Racle-21** | DRB1*15:01 | 0.806 | 242 |
| **Racle-26** | DRB3*02:02 | 0.803 | 356 |
| **Racle-13** | DRB4*01:03 | 0.802 | 287 |
| **Racle-6** | DRB3*02:02 | 0.801 | 207 |
| **Racle-24** | DRB3*02:02 | 0.8 | 256 |
| **Racle-30** | DRB3*02:02 | 0.798 | 435 |
| **Racle-16** | DRB3*01:01 | 0.797 | 264 |
| **Racle-38** | DRB4*01:01 | 0.796 | 218 |
| **Racle-17** | DPA1*01:03-DPB1*02:01 | 0.795 | 326 |
| **Wang-7** | DRB1*01:01 | 0.794 | 119 |
| **Racle-29** | DRB1*03:01 | 0.793 | 226 |
| **Racle-15** | DRB4*01:03 | 0.792 | 493 |
| **Racle-3** | DRB4*01:01 | 0.789 | 354 |
| **Racle-6** | DRB4*01:03 | 0.784 | 216 |
| **Racle-28** | DRB3*02:02 | 0.781 | 254 |
| **Abelin-10** | DRB4*01:03 | 0.778 | 171 |
| **Racle-28** | DRB1*03:01 | 0.775 | 1246 |
| **Racle-1** | DRB4*01:01 | 0.772 | 419 |
| **Racle-29** | DPA1*01:03-DPB1*04:01 | 0.769 | 131 |

| | | | |
|---|---|---|---|
| **Racle-4** | DRB4*01:03 | 0.768 | 259 |
| **Racle-31** | DRB3*01:01 | 0.768 | 543 |
| **Racle-11** | DRB4*01:03 | 0.767 | 445 |
| **Racle-14** | DRB1*04:01 | 0.766 | 138 |
| **Racle-9** | DRB4*01:03 | 0.764 | 651 |
| **Racle-12** | DRB3*02:02 | 0.76 | 204 |
| **Racle-15** | DRB1*03:01 | 0.756 | 2237 |
| **Racle-4** | DRB3*02:02 | 0.754 | 379 |
| **Racle-36** | DPA1*01:03-DPB1*23:01 | 0.75 | 138 |
| **Racle-5** | DPA1*01:03-DPB1*02:01 | 0.749 | 916 |
| **Racle-36** | DRB4*01:03 | 0.748 | 234 |
| **Racle-31** | DPA1*01:03-DPB1*02:01 | 0.744 | 490 |
| **Racle-30** | DRB1*03:01 | 0.743 | 1906 |
| **Racle-44** | DRB4*01:03 | 0.743 | 234 |
| **Racle-3** | DRB1*03:01 | 0.741 | 1737 |
| **Racle-40** | DRB4*01:03 | 0.74 | 112 |
| **Racle-37** | DRB4*01:03 | 0.74 | 508 |
| **Ritz-4** | DRB3*02:02 | 0.738 | 506 |
| **Racle-40** | DRB3*01:01 | 0.737 | 508 |
| **Racle-39** | DRB3*01:01 | 0.737 | 279 |
| **Wang-6** | DRB1*01:01 | 0.737 | 132 |
| **Ritz-2** | DRB1*15:01 | 0.736 | 565 |
| **Racle-42** | DRB4*01:03 | 0.735 | 357 |
| **Racle-46** | DRB3*02:02 | 0.734 | 971 |
| **Wang-1** | DRB1*15:01 | 0.732 | 183 |
| **Racle-33** | DRB3*02:02 | 0.732 | 266 |
| **Saghar-1** | DRB4*01:03 | 0.73 | 112 |
| **Racle-45** | DRB4*01:03 | 0.727 | 147 |
| **Racle-1** | DRB1*03:01 | 0.722 | 1696 |
| **Racle-36** | DPA1*01:03-DPB1*02:01 | 0.721 | 988 |
| **Racle-32** | DRB3*02:02 | 0.712 | 692 |
| **Racle-21** | DRB5*01:01 | 0.711 | 335 |
| **Racle-13** | DRB1*03:01 | 0.709 | 1679 |
| **Racle-36** | DRB5*02:02 | 0.706 | 594 |
| **Racle-46** | DRB1*15:01 | 0.705 | 1188 |
| **Abelin-25** | DRB3*01:01 | 0.704 | 589 |
| **Racle-34** | DRB3*02:02 | 0.704 | 759 |
| **Racle-43** | DPA1*01:03-DPB1*04:01 | 0.703 | 595 |
| **Racle-7** | DRB3*02:02 | 0.702 | 529 |
| **Racle-4** | DPA1*01:03-DPB1*02:01 | 0.7 | 526 |
| **Racle-28** | DPA1*01:03-DPB1*04:01 | 0.7 | 167 |
| **Abelin-1** | DRB4*01:01 | 0.696 | 253 |
| **Racle-35** | DRB4*01:03 | 0.694 | 655 |
| **Abelin-7** | DRB4*01:01 | 0.693 | 196 |

| | | | |
|---|---|---|---|
| **Ritz-2** | DRB1*01:01 | 0.693 | 1278 |
| **Khodadoust-12** | DRB1*01:01 | 0.691 | 198 |
| **Racle-5** | DRB1*11:01 | 0.689 | 407 |
| **Racle-35** | DPA1*01:03-DPB1*23:01 | 0.688 | 157 |
| **Sofron-2** | H-2-IAd | 0.687 | 856 |
| **Racle-18** | DRB1*01:01 | 0.687 | 1504 |
| **Racle-6** | DRB1*11:01 | 0.686 | 767 |
| **Racle-21** | DRB1*07:01 | 0.682 | 294 |
| **Racle-16** | DRB1*01:01 | 0.68 | 1765 |
| **Racle-42** | DPA1*01:03-DPB1*04:01 | 0.678 | 415 |
| **Racle-37** | DRB5*02:02 | 0.677 | 1446 |
| **Racle-12** | DPA1*01:03-DPB1*04:01 | 0.677 | 142 |
| **Abelin-2** | DRB1*03:01 | 0.677 | 1179 |
| **Khodadoust-14** | DRB1*01:01 | 0.675 | 298 |
| **Racle-16** | DPA1*01:03-DPB1*02:01 | 0.675 | 548 |
| **Racle-27** | DRB1*01:01 | 0.674 | 1818 |
| **Racle-41** | DRB1*15:01 | 0.673 | 920 |
| **Ritz-3** | DQA1*01:01-DQB1*05:01 | 0.67 | 405 |
| **Wang-4** | DRB1*15:01 | 0.669 | 165 |
| **Racle-7** | DPA1*01:03-DPB1*02:01 | 0.669 | 705 |
| **Abelin-3** | DRB1*03:01 | 0.669 | 138 |
| **Racle-33** | DPA1*01:03-DPB1*04:01 | 0.667 | 822 |
| **Racle-22** | DRB1*13:01 | 0.667 | 110 |
| **Abelin-8** | DRB4*01:01 | 0.667 | 260 |
| **Racle-35** | DPA1*01:03-DPB1*02:01 | 0.659 | 1082 |
| **Racle-38** | DPA1*01:03-DPB1*04:01 | 0.659 | 137 |
| **Racle-8** | DRB1*01:01 | 0.656 | 1926 |
| **Racle-45** | DRB1*01:01 | 0.656 | 1962 |
| **Racle-46** | DPA1*01:03-DPB1*04:01 | 0.653 | 605 |
| **Racle-19** | DRB1*13:03 | 0.652 | 208 |
| **Racle-36** | DRB1*16:01 | 0.649 | 715 |
| **Racle-26** | DRB1*11:01 | 0.647 | 1616 |
| **Racle-24** | DRB1*11:01 | 0.646 | 1666 |
| **Wang-3** | DRB1*03:01 | 0.643 | 237 |
| **Sofron-1** | H-2-IAd | 0.643 | 1175 |
| **Racle-5** | DPA1*01:03-DPB1*06:01 | 0.642 | 1475 |
| **Racle-9** | DPA1*01:03-DPB1*04:01 | 0.641 | 458 |
| **Racle-32** | DPA1*01:03-DPB1*04:01 | 0.64 | 445 |
| **Racle-24** | DPA1*01:03-DPB1*03:01 | 0.639 | 2158 |
| **Racle-39** | DPA1*01:03-DPB1*02:01 | 0.639 | 228 |
| **Racle-10** | DPA1*01:03-DPB1*04:01 | 0.639 | 973 |
| **Abelin-5** | DRB1*01:01 | 0.638 | 206 |
| **Racle-24** | DRB1*04:05 | 0.636 | 1328 |
| **Sofron-3** | H-2-IAd | 0.636 | 2421 |

| | | | |
|---|---|---|---|
| **Abelin-6** | DRB1*03:01 | 0.636 | 369 |
| **Saghar-2** | DRB3*02:02 | 0.635 | 235 |
| **Racle-28** | DRB1*11:04 | 0.634 | 1855 |
| **Racle-20** | DRB1*07:01 | 0.634 | 402 |
| **Heyder-2** | DRB1*07:01 | 0.63 | 154 |
| **Racle-11** | DRB1*01:03 | 0.629 | 938 |
| **Racle-5** | DRB1*04:04 | 0.629 | 363 |
| **Ooi-2** | DRB1*15:01 | 0.629 | 3327 |
| **Racle-10** | DRB1*04:04 | 0.629 | 402 |
| **Racle-26** | DRB1*04:05 | 0.625 | 2053 |
| **Abelin-9** | DRB4*01:01 | 0.624 | 378 |
| **Racle-29** | DRB1*11:04 | 0.624 | 211 |
| **Racle-2** | DPA1*01:03-DPB1*03:01 | 0.624 | 814 |
| **Ritz-4** | DRB1*01:01 | 0.623 | 2490 |
| **Khodadoust-11** | DRB1*11:01 | 0.622 | 524 |
| **Ritz-2** | DRB5*01:01 | 0.622 | 950 |
| **Racle-4** | DRB1*11:01 | 0.621 | 2930 |
| **Racle-30** | DRB1*11:04 | 0.621 | 2542 |
| **Racle-12** | DRB1*11:01 | 0.621 | 1626 |
| **Racle-45** | DPA1*01:03-DPB1*04:01 | 0.62 | 144 |
| **Abelin-2** | DRB1*01:01 | 0.62 | 1885 |
| **Abelin-4** | DRB1*11:01 | 0.619 | 132 |
| **Racle-45** | DPA1*01:03-DPB1*02:01 | 0.618 | 1272 |
| **Abelin-13** | DRB1*03:01 | 0.618 | 1006 |
| **Racle-25** | DPA1*01:03-DPB1*03:01 | 0.617 | 2484 |
| **Racle-35** | DRB5*02:02 | 0.616 | 1497 |
| **Racle-43** | DRB1*04:01 | 0.616 | 967 |
| **Racle-25** | DRB1*11:01 | 0.615 | 422 |
| **Racle-6** | DRB1*04:04 | 0.613 | 1482 |
| **Racle-4** | DPA1*01:03-DPB1*06:01 | 0.612 | 1445 |
| **Wang-2** | DRB1*11:01 | 0.61 | 117 |
| **Wang-1** | DRB1*03:01 | 0.609 | 225 |
| **Racle-15** | DRB1*04:01 | 0.609 | 2529 |
| **Racle-13** | DPA1*01:03-DPB1*04:01 | 0.608 | 242 |
| **Racle-41** | DPA1*01:03-DPB1*04:01 | 0.608 | 216 |
| **Ooi-1** | DRB1*15:01 | 0.607 | 1495 |
| **Racle-27** | DPA1*01:03-DPB1*06:01 | 0.606 | 723 |
| **Racle-17** | DRB1*13:01 | 0.604 | 317 |
| **Wang-7** | DRB1*04:01 | 0.603 | 216 |
| **Sofron-4** | H-2-IAb | 0.603 | 734 |
| **Racle-25** | DRB1*04:05 | 0.602 | 288 |
| **Abelin-13** | DRB1*01:01 | 0.602 | 1226 |
| **Racle-20** | DRB1*13:03 | 0.6 | 701 |
| **Abelin-12** | DRB1*04:01 | 0.597 | 1138 |

| | | | |
|---|---|---|---|
| **Sofron-5** | H-2-IAb | 0.596 | 2493 |
| **Heyder-2** | DRB1*04:01 | 0.595 | 516 |
| **Racle-46** | DPA1*01:03-DPB1*03:01 | 0.594 | 3339 |
| **Abelin-11** | DRB5*01:01 | 0.593 | 301 |
| **Abelin-1** | DRB1*04:05 | 0.592 | 2135 |
| **Racle-8** | DPA1*01:03-DPB1*20:01 | 0.592 | 1867 |
| **Racle-13** | DRB1*04:01 | 0.59 | 2082 |
| **Racle-4** | DRB1*04:04 | 0.59 | 2302 |
| **Racle-11** | DRB1*04:04 | 0.588 | 1747 |
| **Racle-9** | DRB1*01:03 | 0.586 | 1547 |
| **Abelin-11** | DRB1*04:01 | 0.586 | 1345 |
| **Racle-37** | DRB1*16:01 | 0.585 | 1736 |
| **Racle-3** | DRB1*07:01 | 0.584 | 1536 |
| **Racle-41** | DQA1*01:01-DQB1*05:01 | 0.584 | 113 |
| **Racle-36** | DRB1*07:01 | 0.579 | 470 |
| **Racle-9** | DRB1*04:04 | 0.578 | 2543 |
| **Racle-1** | DPA1*01:03-DPB1*03:01 | 0.576 | 1781 |
| **Racle-10** | DRB1*01:03 | 0.573 | 297 |
| **Abelin-3** | DRB1*11:01 | 0.572 | 162 |
| **Racle-22** | DRB1*14:01 | 0.565 | 103 |
| **Abelin-4** | DRB1*03:01 | 0.565 | 121 |
| **Racle-46** | DRB5*01:01 | 0.564 | 1050 |
| **Nanaware-2** | DRB1*01:01 | 0.564 | 2455 |
| **Racle-8** | DPA1*01:03-DPB1*04:01 | 0.564 | 105 |
| **Abelin-7** | DRB1*07:01 | 0.563 | 346 |
| **Racle-41** | DRB5*01:01 | 0.559 | 2496 |
| **Abelin-8** | DRB1*07:01 | 0.558 | 405 |
| **Racle-35** | DRB1*07:01 | 0.558 | 991 |
| **Wang-5** | DRB1*04:01 | 0.557 | 846 |
| **Wang-6** | DRB1*04:01 | 0.557 | 156 |
| **Ooi-4** | DRB1*01:01 | 0.556 | 3986 |
| **Racle-33** | DRB1*11:01 | 0.554 | 4209 |
| **Khodadoust-6** | DRB1*01:01 | 0.551 | 240 |
| **Racle-41** | DRB1*01:02 | 0.549 | 4126 |
| **Racle-35** | DRB1*16:01 | 0.548 | 2296 |
| **Racle-44** | DRB1*04:01 | 0.547 | 1943 |
| **Heyder-1** | DRB1*04:01 | 0.547 | 388 |
| **Racle-42** | DRB1*04:01 | 0.545 | 3028 |
| **Abelin-5** | DRB1*11:01 | 0.544 | 227 |
| **Abelin-12** | DRB5*01:01 | 0.544 | 264 |
| **Heyder-3** | DRB1*03:01 | 0.543 | 193 |
| **Racle-45** | DRB1*04:08 | 0.542 | 2892 |
| **Racle-23** | DRB1*13:01 | 0.54 | 780 |
| **Clement-1** | DRB1*01:01 | 0.538 | 2963 |

| | | | |
|---|---|---|---|
| **Racle-37** | DRB1*07:01 | 0.534 | 2061 |
| **Wang-5** | DRB1*10:01 | 0.534 | 925 |
| **Khodadoust-6** | DRB1*04:01 | 0.531 | 680 |
| **Saghar-4** | DRB1*01:02 | 0.528 | 568 |
| **Racle-40** | DRB1*04:04 | 0.527 | 1809 |
| **Racle-1** | DRB1*07:01 | 0.527 | 818 |
| **Nanaware-1** | DRB1*01:01 | 0.526 | 3596 |
| **Racle-32** | DRB1*11:01 | 0.521 | 6101 |
| **Racle-34** | DRB1*11:01 | 0.518 | 6633 |
| **Racle-39** | DRB1*04:04 | 0.514 | 2706 |
| **Wang-4** | DRB1*04:03 | 0.511 | 516 |
| **Khodadoust-4** | DRB1*04:01 | 0.507 | 314 |
| **Racle-45** | DQA1*01:01-DQB1*05:01 | 0.504 | 131 |
| **Nanaware-3** | DRB1*01:01 | 0.504 | 4418 |
| **Abelin-23** | DRB1*15:01 | 0.503 | 2598 |
| **Nanaware-4** | DRB1*01:01 | 0.503 | 3207 |
| **Abelin-21** | DRB1*12:01 | 0.503 | 854 |
| **Khodadoust-12** | DRB1*07:01 | 0.502 | 255 |
| **Racle-7** | DRB1*11:04 | 0.502 | 5565 |
| **Abelin-9** | DRB1*07:01 | 0.501 | 842 |
| **Racle-24** | DQA1*05:05-DQB1*03:01 | 0.5 | 123 |
| **Racle-18** | DRB1*13:01 | 0.499 | 1333 |
| **Racle-38** | DPA1*01:03-DPB1*03:01 | 0.499 | 1039 |
| **Ting-1** | DRB1*04:05 | 0.499 | 1665 |
| **Khodadoust-14** | DRB1*07:01 | 0.497 | 486 |
| **Benito-1** | DRB1*03:01 | 0.492 | 1285 |
| **Scally-1** | DRB1*04:01 | 0.492 | 771 |
| **Racle-38** | DRB1*07:01 | 0.49 | 615 |
| **Racle-44** | DRB1*08:01 | 0.489 | 2420 |
| **Racle-31** | DQA1*01:10-DQB1*06:03 | 0.486 | 122 |
| **Ritz-1** | DQA1*01:02-DQB1*06:02 | 0.486 | 346 |
| **Abelin-14** | DPA1*01:03-DPB1*06:01 | 0.485 | 1795 |
| **Abelin-11** | DRB1*15:03 | 0.485 | 184 |
| **Karunakaran-1** | H-2-IAb | 0.484 | 1544 |
| **Racle-23** | DRB1*14:01 | 0.484 | 354 |
| **Racle-45** | DQA1*03:03-DQB1*03:01 | 0.484 | 141 |
| **Racle-26** | DPA1*01:03-DPB1*03:01 | 0.484 | 104 |
| **Abelin-26** | DPA1*01:03-DPB1*02:01 | 0.482 | 2164 |
| **Racle-36** | DQA1*01:02-DQB1*05:02 | 0.476 | 206 |
| **Abelin-1** | DRB1*07:01 | 0.472 | 1454 |
| **Racle-8** | DRB4*01:03 | 0.469 | 107 |
| **Scally-3** | DRB1*04:04 | 0.463 | 1809 |
| **Abelin-15** | DQA1*01:02-DQB1*06:04 | 0.46 | 981 |
| **Ooi-1** | DRB5*01:01 | 0.458 | 930 |

| | | | |
|---|---|---|---|
| **Racle-16** | DRB1*13:01 | 0.457 | 1825 |
| **Racle-46** | DRB1*12:01 | 0.456 | 922 |
| **Nelde-3** | DRB1*07:01 | 0.455 | 1372 |
| **Racle-46** | DQA1*01:02-DQB1*05:02 | 0.453 | 238 |
| **Abelin-17** | DRB1*03:01 | 0.451 | 2118 |
| **Racle-31** | DRB1*10:01 | 0.444 | 1797 |
| **Abelin-10** | DRB1*04:01 | 0.438 | 107 |
| **Ritz-4** | DRB1*13:01 | 0.435 | 2561 |
| **Ritz-3** | DQA1*01:03-DQB1*06:03 | 0.429 | 1607 |
| **Racle-37** | DQA1*01:02-DQB1*05:02 | 0.427 | 123 |
| **Khodadoust-10** | DRB1*04:01 | 0.426 | 814 |
| **Ooi-3** | DRB1*01:01 | 0.424 | 4254 |
| **Racle-42** | DRB1*08:01 | 0.422 | 5777 |
| **Scally-2** | DRB1*04:02 | 0.421 | 1083 |
| **Abelin-12** | DRB1*15:03 | 0.419 | 152 |
| **Racle-8** | DRB1*07:01 | 0.417 | 424 |
| **Racle-38** | DQA1*02:01-DQB1*02:02 | 0.416 | 697 |
| **Racle-43** | DRB1*08:01 | 0.416 | 2348 |
| **Racle-25** | DQA1*05:05-DQB1*03:01 | 0.412 | 273 |
| **Nanaware-5** | H-2-IAb | 0.407 | 481 |
| **Khodadoust-5** | DRB1*04:01 | 0.406 | 672 |
| **Racle-39** | DQA1*01:03-DQB1*06:03 | 0.402 | 318 |
| **Racle-31** | DRB1*13:01 | 0.399 | 2273 |
| **Racle-40** | DRB1*13:01 | 0.398 | 1568 |
| **Racle-35** | DQA1*01:02-DQB1*05:02 | 0.397 | 336 |
| **Abelin-27** | DQA1*01:02-DQB1*06:02 | 0.389 | 2950 |
| **Racle-46** | DQA1*05:05-DQB1*03:01 | 0.388 | 338 |
| **Racle-1** | DQA1*02:01-DQB1*02:01 | 0.388 | 293 |
| **Racle-37** | DQA1*02:01-DQB1*02:02 | 0.385 | 165 |
| **Abelin-16** | DRB1*01:01 | 0.384 | 6298 |
| **Racle-34** | DQA1*05:05-DQB1*03:01 | 0.382 | 410 |
| **Abelin-20** | DRB1*11:01 | 0.381 | 7588 |
| **Racle-38** | DRB1*13:01 | 0.381 | 2778 |
| **Saghar-3** | DRB1*16:01 | 0.38 | 1385 |
| **Abelin-19** | DRB1*07:01 | 0.378 | 4297 |
| **Racle-40** | DQA1*01:03-DQB1*06:03 | 0.377 | 181 |
| **Khodadoust-9** | DRB1*11:01 | 0.376 | 1218 |
| **Racle-33** | DQA1*05:05-DQB1*03:01 | 0.375 | 937 |
| **Racle-36** | DQA1*02:01-DQB1*02:02 | 0.375 | 964 |
| **Racle-42** | DQA1*04:01-DQB1*03:01 | 0.372 | 619 |
| **Racle-4** | DQA1*05:05-DQB1*03:01 | 0.363 | 203 |
| **Racle-39** | DRB1*13:01 | 0.363 | 1174 |
| **Khodadoust-8** | DRB1*04:01 | 0.359 | 628 |
| **Racle-21** | DPA1*01:03-DPB1*17:01 | 0.359 | 425 |

| | | | |
|---|---|---|---|
| **Racle-43** | DQA1*04:01-DQB1*03:01 | 0.358 | 726 |
| **Racle-44** | DQA1*04:01-DQB1*03:01 | 0.358 | 134 |
| **Khodadoust-13** | DRB1*01:01 | 0.354 | 986 |
| **Racle-5** | DQA1*05:05-DQB1*03:01 | 0.353 | 224 |
| **Khodadoust-15** | DRB1*07:01 | 0.352 | 373 |
| **Racle-35** | DQA1*02:01-DQB1*02:02 | 0.35 | 987 |
| **Racle-32** | DQA1*05:05-DQB1*03:01 | 0.347 | 923 |
| **Khodadoust-1** | DRB1*04:01 | 0.343 | 3677 |
| **Racle-10** | DQA1*05:05-DQB1*03:01 | 0.343 | 366 |
| **Wang-2** | DRB1*13:02 | 0.333 | 140 |
| **Nelde-3** | DRB1*08:01 | 0.333 | 1844 |
| **Abelin-22** | DRB1*13:03 | 0.33 | 2677 |
| **Khodadoust-11** | DRB1*07:01 | 0.329 | 649 |
| **Abelin-10** | DRB5*01:01 | 0.328 | 221 |
| **Abelin-18** | DRB1*04:01 | 0.326 | 1960 |
| **Saghar-2** | DRB1*14:01 | 0.323 | 1994 |
| **Saghar-1** | DRB1*09:01 | 0.321 | 843 |
| **Abelin-24** | DRB1*16:01 | 0.318 | 919 |
| **Racle-13** | DQA1*05:01-DQB1*02:01 | 0.315 | 166 |
| **Nelde-1** | DRB1*08:03 | 0.315 | 1013 |
| **Racle-41** | DQA1*01:02-DQB1*06:02 | 0.312 | 409 |
| **Racle-25** | DQA1*05:05-DQB1*02:02 | 0.306 | 189 |
| **Khodadoust-1** | DRB1*07:01 | 0.302 | 655 |
| **Racle-12** | DQA1*05:05-DQB1*03:01 | 0.295 | 163 |
| **Racle-9** | DQA1*05:05-DQB1*03:01 | 0.295 | 166 |
| **Khodadoust-3** | DRB1*01:01 | 0.288 | 528 |
| **Bergseng-3** | DQA1*05:01-DQB1*03:01 | 0.281 | 6149 |
| **Racle-16** | DQA1*01:03-DQB1*06:03 | 0.277 | 125 |
| **Khodadoust-7** | DRB1*07:01 | 0.272 | 1095 |
| **Racle-7** | DQA1*05:05-DQB1*03:01 | 0.27 | 395 |
| **Nelde-2** | DRB1*13:01 | 0.263 | 190 |
| **Khodadoust-2** | DRB1*11:01 | 0.26 | 3219 |
| **Bergseng-1** | DQA1*02:01-DQB1*02:02 | 0.233 | 5823 |
| **Bergseng-2** | DQA1*05:01-DQB1*02:01 | 0.224 | 3052 |
| **Khodadoust-2** | DRB1*07:01 | 0.217 | 1512 |
| **Khodadoust-3** | DRB1*04:01 | 0.211 | 1397 |
| **Sofron-3** | H-2-IEd | 0.204 | 306 |
| **Nelde-2** | DRB1*08:01 | 0.201 | 177 |
| **Nelde-1** | DRB1*13:01 | 0.188 | 2066 |
| **Racle-41** | DPA1*01:03-DPB1*05:01 | 0.186 | 591 |
